# MinC influences the Min oscillation period in *Escherichia coli* by priming MinD on the membrane

**DOI:** 10.64898/2026.10.05.756812

**Authors:** Colby N. Ferreira, Catherine E. Trebino, Samuel Bartlett, Joseph Conti, Christopher J. LaBreck, Jodi L. Camberg

## Abstract

Spatial organization during bacterial fission is critical to ensure that division results in identical, viable progeny cells. In *Escherichia coli,* one important system that accomplishes this is the Min system, comprising MinC, MinD, and MinE proteins. The Min system regulates assembly of the FtsZ-ring to ensure proper placement of the division machinery at midcell. We explored the dynamics of the Min system and demonstrate that Min oscillation and release of MinD from the membrane is modulated by MinC. Deletion of the *minC* gene from a strain expressing *minD* fused to the gene encoding green fluorescent protein (Gfp) resulted in faster pole-to-pole oscillation of Gfp-MinD in vivo, leading to a shorter period. This was also observed in a strain expressing the MinC(R133A), a variant mutated at the MinC-MinD interface. In phospholipid binding and release assays in vitro, we observed that the concentration of MinE required to release MinD was lower with MinC, suggesting that MinC activates MinD for MinE-dependent release. Oscillation is produced by a diffusion-capture-release mechanism occurring at the membrane with proteins proceeding toward a pole. To determine if an increase in the diffusive proportion of MinD leads to faster oscillation, we constructed MinD membrane targeting sequence (MTS) variants to reduce the membrane associated pool of MinD. MTS variants induced a range of oscillation defects, with MinD(KR265AA) oscillating faster in vivo. Together our results show that MinC is an effector that modifies Min oscillation by priming MinD for MinE-dependent membrane release and fine-tunes the Min oscillation rate in dividing cells.

**Importance:** Gram negative bacteria, such as *Escherichia coli,* divide at the longitudinal center, giving rise to identical progeny cells. This requires polymerization of FtsZ into a ring, and FtsZ-ring placement is controlled by Min oscillation between the cell poles. Here we show that the FtsZ inhibitor MinC has a surprising capacity to regulate the Min oscillation rate through modifying the membrane release step of MinD.

## Introduction

Prokaryotic cell division begins with the assembly and regulation of a large cytoskeletal protein structure known as the FtsZ-ring, or ‘Z-ring’. In all bacteria, assembly of the FtsZ-ring is the hallmark of division (1, 2). This ring is comprised primarily of FtsZ and FtsZ-interacting proteins. In *E. coli,* these proteins include FtsA and ZipA, which tether the ring to the cytoplasmic membrane (3–5). After assembly of the FtsZ-ring, additional proteins are recruited to the division site, such as FtsN, FtsI, and FtsW, leading to peptidoglycan remodeling and septal cell wall synthesis (6). Mislocalization of the FtsZ-ring leads to cell division defects indicated by the presence of long filamentous cells and small minicells (7–9).

Selection of the division plane in *E. coli* depends on two mechanisms: (i) nucleoid occlusion by SlmA, which functions by binding to DNA and FtsZ preventing formation of the FtsZ-ring over the chromosomes, and (ii) the Min system, which functions by directly inhibiting FtsZ polymerization at the poles (10–12). *E. coli* lacking both of these systems are non-viable (13). It has also been shown that in the pathogenic *E. coli* strains 0157:H7 and *Neisseria gonorrhea*, deletion of the Min system leads to reduced virulence (14, 15). *E. coli* cells lacking the Min system display filamentation and minicells that arise from aberrant division events. *N. gonorrhoeae* cells deleted for *min* genes display enlarged cells and a high frequency of lysis (14). Additionally, it has been observed that *Helicobacter pylori* cells deleted for *minC* display decreased motility, growth defects, and altered morphology, implicating MinC in cell shape development (16). The Min system in *E. coli* includes three proteins, MinC, MinD, and MinE. MinD is an ATPase that cycles through steps of ATP binding and hydrolysis, which direct membrane engagement, and this is regulated by MinE (17–19). Inside the cell, this collective activity produces movement of the Min proteins between the cell poles, termed oscillation (20, 21).

MinC binds to MinD, sharing a partially overlapping binding site with MinE on MinD, and MinC is a passenger to MinD oscillation (19). MinC functions to destabilize FtsZ polymers and prevent ring assembly near the poles (8, 11). MinC binds to the FtsZ-FtsZ subunit interface preventing subunits from assembling head to tail into the FtsZ polymer and to an extended FtsZ C-terminal region (10, 11). The current model suggests that MinD binds to the cytoplasmic membrane in the presence of ATP, and MinD is stimulated for ATP hydrolysis and release from the membrane by MinE, orchestrating the cycle of membrane binding and release that produces oscillation (17, 20–22). Only MinD and MinE are required to establish oscillation in vitro and in vivo, leading to the suggestion that oscillation is unaltered by MinC.

MinC is dimeric and each protomer is comprised of two separate domains, the N- and C-domains, connected by a short linker (22). The C-terminal domain is a beta-helix and supports dimerization. The C-terminal domain of MinC is the site of MinD binding, with the interface consisting of residues that include R94, A95, D110, N112, K128, and R130 of MinC of *Aquifex aeolicus* (23). In *E. coli* several of these contacts are predicted to be conserved, with R133, which corresponds to R94 in

*A. aeolicus*, likely forming a salt bridge with residue D154 of MinD (24). The C-domain of MinC also binds the extended C-terminal region of FtsZ and is thought to prevent lateral bundling of FtsZ filaments (11). The MinC N-terminal domain consists of two alpha-helices and 5 beta-strands, which are thought bind to the FtsZ-FtsZ subunit interface and prevent head to tail assembly of FtsZ protomers (22). The amino acid, G10, on the N-terminus of MinC has been reported to be important for FtsZ disassembly (8). Our group and others mapped additional N-domain residues to a binding cleft, including K9, G10, S12, S16, K35, A39, and F42 on MinC that are important for FtsZ interaction and polymer disassembly (8, 11).

MinD, the energy-harnessing component of the oscillatory system, is a member of the Walker A cytoskeletal ATPase (WACA) protein family and contains a deviant Walker A motif with a signature lysine that plays a role in hydrolysis and dimerization (9, 17, 18, 24, 25). MinD associates with the inner cytoplasmic membrane through a C-terminal amphipathic helix known as its membrane targeting sequence (MTS) (17, 18). Stable membrane association by MinD occurs in the ATP bound dimer conformation. MinE then binds to MinD at the membrane and stimulates ATP hydrolysis, promoting releasing MinD from the membrane surface (9, 19, 22). MinE exists in two distinct conformations, a 6-β-stranded “closed” conformation, and a 4-β-stranded “open conformation”; the “open conformation” is capable of binding to MinD (26–28). After removal of MinD from the membrane, MinE lingers at the membrane, anchored by its N-terminal membrane targeting helix, presumably preventing MinD from rebinding at that position (17–19). This is supported by in vitro reconstitutions where a leading edge of MinD is followed by a lagging edge of MinE, as observed on lipid monolayers (29–31).

Our group demonstrated that MinC and MinD dimers assemble into copolymers in vitro in the presence of ATP (27). These copolymers were suggested to be alternating dimers of MinC and MinD by the Lowe group, as detected in the crystal structure of Min proteins from *Aquifex aeolicus* (23). Upon addition of ATP, MinC-MinD polymers form rapidly and remain stable for long periods. Addition of MinE to the reaction, or the protease complex ClpXP, destabilizes the MinC-MinD copolymers and promotes disassembly (32, 33). MinC and MinD proteins of *Pseudomonas aeruginosa* and *Bacillus subtilis* have also exhibited polymers (34, 35). Although MinC-MinD copolymers form readily in vitro and have been observed by electron microscopy, they are likely short lived in vivo due to the presence of MinE, which stimulates MinD to cycle through ATP hydrolysis steps. MinC is also not required for robust oscillation in vivo. Cells expressing a heterodimer of MinD that fails to form copolymers, but still binds MinC, confers wild-type morphology (7). Thus, MinC-MinD copolymer formation is not required for oscillation in vivo, and it remains unclear how stable they are in vivo.

Here to determine if MinC has the capacity to modulate oscillation of the Min system as an effector, we tested if mutation of amino acid residues in the MinC-MinD interface confers Min oscillation defects in dividing cells. MinD oscillation occurs through cycles of membrane attachment and release, regulated by MinE, and the current paradigm assumes that MinC does not impact oscillation (9, 29). However, our group previously observed that mutations in MinC, including S16D and L194P, altered the Min oscillation rate in vivo, suggesting that MinC plays a more important role in oscillation than cargo (8). Using strains engineered to express *gfp-minD* or *gfp-minC* from the chromosome, combined with direct binding and recruitment assays in vitro, our results show that MinC functions as an effector of the Min system to modify MinE-stimulated release of MinD from the membrane and regulate oscillation timing in the cell.

## Results

### Mutations in *minC* alter redistribution timing of Gfp-MinD in dividing cells

To determine if the presence or absence of MinC contributes to the timing of Min oscillation, we compared MinD pole-to-pole oscillation in dividing cells with and without *minC*. First, we constructed a strain to express Gfp-MinD from the native locus by replacing the chromosomal copy of *minD* with *gfp-minD* in the *min* operon by site-specific recombineering (CF005) (Table 1). Expression of Gfp-MinD from the chromosome caused no observable morphological defects under the conditions tested and did not induce cell filamentation (Fig. 1Ai) (Fig. S1A). The mean length of cells expressing Gfp-MinD was identical to wild type MG1655 cells, 1.9 +/- 0.03 µm and 1.9 +/- 0.03 µm, respectively, and no minicells were present (Fig. S1B) (Table 2).

**Fig. 1.**
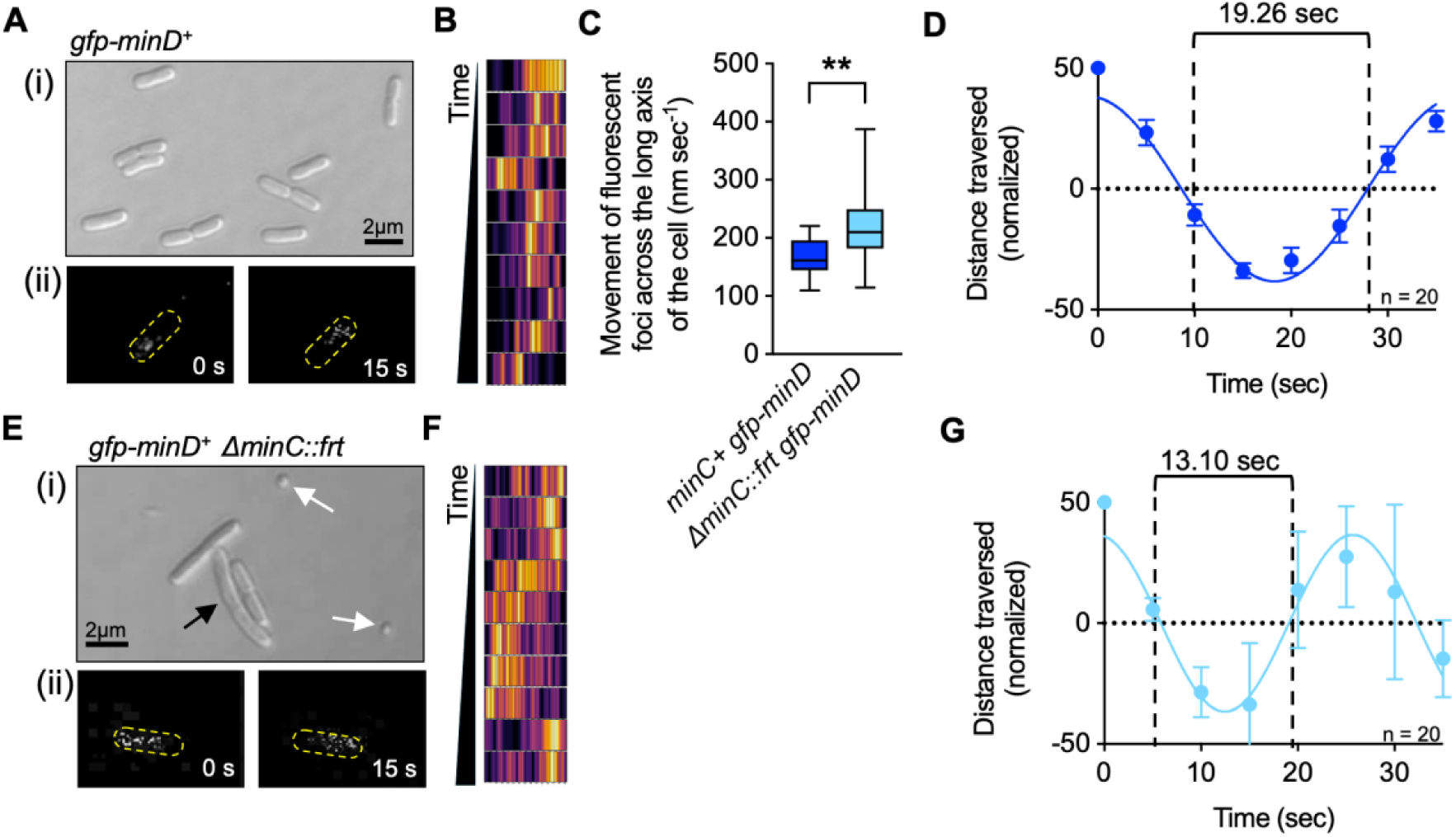
Deletion of *minC* from a strain expressing *gfp-minD* from the chromosome results in modified Min oscillation. (A) Morphology and Gfp-MinD oscillation of a strain expressing *gfp-minD* (CF005) were observed by (i) DIC microscopy and (ii) confocal fluorescence microscopy at 5 second time intervals. Scale bar is 2 µm. (B) Kymograph of Gfp-MinD movement across a single live cell shown as a 50 second time series with images taken every 5 seconds. (C) Oscillation rate of Gfp-MinD was observed in the presence or absence of chromosomal MinC (CF005 and CF315, respectively). Rates are calculated from at least 20 cells. (D) A plot of the periodic wave formed by Gfp-MinD across the cell in the presence of MinC is shown (CF005). Data from 20 cells was fit to a sine wave. (E) Morphology and Gfp-MinD oscillation of a strain deleted for *minC* (CF315) by (i) DIC microscopy, and (ii) confocal fluorescence microscopy. Black arrows denote filamentous cells while white arrows denote minicells. Scale bar is 2 µm. (F) Kymograph of Gfp-MinD movement across a live cell, in the absence of MinC, during a 50 second time series with time lapse images taken every 5 seconds. (G) A plot of the periodic wave formed by Gfp-MinD across the cell in a strain deleted for *minC* (CF315) is shown. Data from 20 cells was fit to a sine wave. In (C), *p* values are as follows, ∗∗ < 0.005. Error is shown as standard error.

**Table 1.** *E. coli* strains.

| Strain or Plasmid | Relevant Genotype | Source, reference, or construction |
| --- | --- | --- |
| MG1655 | lambda<sup>-</sup> *rph-1* | (48) |
| BL21(DE3) Δ*clpP* | *F-ompT gal dcm lon hsdSB(rB- mB-) $\lambda$(DE3)[lacI lacUV5-T7 gene 1 ind1 sam7 nin5]) clpP::cat* | S. Wickner<sup>1</sup> |
| CL0030 | MG1655 Δ*minC::kan-P<sub>rha</sub>-parE* | MG1655; λred |
| CL0248 | MG1655 Δ*minC::P<sub>BAD</sub>-gfp-minC* | CL0030; MG1655; λred |
| CL0481 | MG1655 Δ*minC::P<sub>BAD</sub>-gfp-minC ΔminD::kan-Prha-parE* | CL0248; λred |
| CL0476 | MG1655 Δ*minD::kan-P<sub>rha</sub>-parE* | MG1655; λred |
| CF005 | MG1655 Δ*minD::gfp-minD* | CL0476; λred |
| CF010 | MG1655 Δ*minD::gfp-minD ΔminC::kan-P<sub>rha</sub>-parE* | CF005; λred |
| CF254 | MG1655 Δ*minC::P<sub>BAD</sub>-gfp-minC ΔminD::minD(FL263GG)* | CL0481; λred |
| CF071 | MG1655 Δ*minC::P<sub>BAD</sub>-gfp-minC ΔminD::minD(KR265AA)* | CL0481; λred |
| CL0482 | MG1655 Δ*minC::P<sub>BAD</sub>-gfp-minC ΔminD::minD(KR265EE)* | CL0481; λred |
| CF070 | MG1655 Δ*minC::P<sub>BAD</sub>-gfp-minC ΔminD::minD(Δ8)* | CL0481; λred |
| CF086 | MG1655 Δ*minC::P<sub>BAD</sub>-gfp-minC ΔminD::minD(D154A)* | CL0481; λred |
| CF266 | MG1655 Δ*minC::P<sub>BAD</sub>-gfp-minC(R133A)* | CL030; λred |
| CF150 | MG1655 Δ*minD::gfp-minD(FL263GG)* | CL0476; λred |
| CF097 | MG1655 Δ*minD::gfp-minD(KR265AA)* | CL0476; λred |
| CF255 | MG1655 Δ*minD::gfp-minD(KR265EE)* | CL0476; λred |
| CF256 | MG1655 Δ*minD::gfp-minD(Δ8)* | CL0476; λred |
| CF089 | MG1655 Δ*minD::gfp-minD(D154A)* | CL0476; λred |
| CF0312 | MG1655 Δ*minD::gfp-minD(D154A) ΔminC::cat* | CF089; λred |
| CF0313 | MG1655 Δ*minD::gfp-minD(D154A) ΔminC::frt* | CF0312; pCP20 |
| CF095 | MG1655 Δ*minD::gfp-minD ΔminC::minC(R133A)* | CF010; λred |
| CF0314 | MG1655 Δ*minD::gfp-minD ΔminC::cat* | CF005; λred |
| CF0315 | MG1655 Δ*minD::gfp-minD ΔminC::frt* | CF0314; pCP20 |

<sup>1</sup> BL21(DE3) $\Delta$ *clpP* was a gift from Sue Wickner

**Table 2.** Morphology of Engineered MinD and MinC strains.

| Genotype | Cell Length | Minicell Percentage | Oscillation Rate |
| --- | --- | --- | --- |
| MG1655 | 1.9 ± 0.03 mm | 0% | n.d.<sup>1</sup> |
| ***gfp-minD background*** |  |  |  |
| *minC gfp-minD* | 1.9 ± 0.03 mm | 0% | 166.0 ± 7.3 nm sec<sup>-1</sup> |
| *ΔminC::frt gfp-minD* | 3.9 ± 0.15 mm | 10.31% | 216.9 ± 12.7 nm sec<sup>-1</sup> |
| *minC(R133A) gfp-minD* | 3.5 ± 0.16 mm | 8.67% | 218.8 ± 22.7 nm sec<sup>-1</sup> |
| *minC gfp-minD(D154A)* | 3.1 ± 0.10 mm | 6.97% | 184.3 ± 6.0 nm sec<sup>-1</sup> |
| *ΔminC::frt gfp-minD(D154A)* | 3.3 ± 0.13 mm | 7.23% | 186.2 ± 4.2 nm sec<sup>-1</sup> |
| *minC gfp-minD(KR265AA)* | 2.6 ± 0.08 mm | 7.04% | 185.7 ± 9.0 nm sec<sup>-1</sup> |
| *minC gfp-minD(Δ8)* | 3.3 ± 0.10 mm | 6.10% | n.d. |
| *minC gfp-minD(FL263GG)* | 2.7 ± 0.15 mm | 12.22% | n.d. |
| *minC gfp-minD(KR265EE)* | 2.9 ± 0.09 mm | 2.91% | n.d. |
| ***gfp-minC background*** |  |  |  |
| *gfp-minC minD* | 1.6 ± 0.09 mm | 0% | 165.4 ± 5.1 nm sec<sup>-1</sup> |
| *gfp-minC(R133A) minD* | 3.5 ± 2.34 mm | 10.71% | n.d. |
| *gfp-minC minD(D154A)* | 2.9 ± 0.12 mm | 10.31% | 188.8 ± 4.8 nm sec<sup>-1</sup> |
| *gfp-minC ΔminD::kan* | 1.6 ± 0.64 mm | 9.31% | n.d. |
| *gfp-minC minD(KR265AA)* | 2.4 ± 0.08 mm | 11.11% | 191.8 ± 11.1 nm sec<sup>-1</sup> |
| *gfp-minC minD(Δ8)* | 3.2 ± 0.09 mm | 6.97% | n.d. |
| *gfp-minC minD(FL263GG)* | 2.4 ± 0.09 mm | 7.40% | n.d. |
| *gfp-minC minD(KR265EE)* | 3.0 ± 0.07 mm | 3.84% | n.d. |

<sup>1</sup> n.d. indicates oscillation was not observed.

Then, we monitored pole-to-pole oscillation of Gfp-MinD in live dividing cells, which we collected from log phase cultures, by confocal fluorescence microscopy (Fig. 1Aii). In single cells, fluorescence was imaged at five second intervals to collect data over the course of 60 seconds to measure the oscillation period (Fig. 1Aii). At every time point, we mapped the fluorescent foci relative to the distance traversed in each cell (Fig. 1B). The time (t) to traverse a cell length (*d*) was used to calculate the rate (*d*×t^-1^), and the rates from 20 cells were averaged. This rate reflects the movement of oscillation and reports the bulk reaction-diffusion velocity, which we calculated to be 166.0 +/- 7.3 nm sec^-1^ for Gfp-MinD in a live cell (Fig. 1C). This measurement describes the movement in a manner that incorporates each cell’s length, instead of assuming a fixed cell length across a population. We also modeled the data as a sine wave to determine the average period for a complete oscillation cycle for the population. We found that in our parental background (MG1655), the complete oscillation cycle of Gfp-MinD is 38.52 sec, and it takes 19.26 sec to move from the middle of a cell, to a pole, and then back to the middle (Fig. 1D).

Next, to test if the absence of MinC changes the bulk reaction-diffusion velocity that we observed for Gfp-MinD, we deleted *minC* by replacing it with, and then excising, an antibiotic resistance gene (*cat)*. This left a short *frt* scar but allowed read-through of the *min* operon to *gfp-minD*, which we confirmed by visualizing Gfp-MinD expression by immunoblot (Fig S1C). As expected, cells deleted for *minC* were mildly filamentous, with an average cell length of 3.9 ± 0.15 µm (Fig. 1Ei) (Fig S1A and S1B) (Table 2). We also observed a high percentage of minicells (10.31%) (Fig. 1Ei) (Fig S1A) (Table 2), consistent with previous reports of strains deleted for *minC* (33). In cells without *minC* (CF315), Gfp-MinD oscillated from pole-to-pole as visualized by confocal fluorescence microscopy (Fig. 1Eii, Fig. 1F); however, when we measured the bulk reaction-diffusion velocity, it was 48% faster in cells without *minC* than in cells containing *minC* (216.9 ± 12.7 nm sec^-1^, and 166.0 ± 7.3 nm sec^-1^ respectively) (Fig. 1C). This suggests that although MinC is not required for oscillation, its presence in cells leads to slower redeployment of Gfp-MinD from a pole across the long axis of the cell. This implies that MinC functions to regulate the oscillation of Gfp-MinD. Accordingly, modeling the oscillation as a sine wave indicated that without MinC, the complete oscillation period of Gfp-MinD to travel from one pole, to the opposite pole, and then return, is 32% shorter in cells without *minC* than in cells with *minC* (26.20 sec and 38.52 seconds, respectively), because Gfp-MinD relocalizes faster across the cell via bulk diffusion (Fig 1D and Fig 1G).

It was previously reported that amino acid substitutions mapping to the MinC-MinD interface prevent MinC from binding to MinD, and it was further shown that mutations at this interface do not prevent MinD oscillation; these substitution mutations include R133A of MinC and D154A of MinD (Fig. 2A) (7, 33). Therefore, to test if cells expressing the MinC mutant protein MinC(R133A), which is defective for binding to MinD, also exhibit fast Gfp-MinD oscillation, we replaced chromosomal *minC* with *minC(R133A)* in cells expressing Gfp-MinD, by using an antibiotic resistance gene linked to *parE* for secondary negative selection, and then measured the reaction-diffusion velocity in live cells (CF095). We observed faster Gfp-MinD oscillation in cells expressing MinC(R133A), 218.8 ± 22.7 nm sec^-1^ (n= 20), than in cells with wild type *minC* (Fig. 2B) (Fig. S1D), similar to the rate observed in cells deleted for *minC*. In addition, cells containing MinC(R133A) have an average cell length of 3.5 ± 0.16 µm, and we observed the presence of short filaments and minicells (8.67%) (Fig. S1A) (Table 2), similar to the morphology of cells deleted for *minC*, suggesting that MinC(R133A) is defective for function. These results indicate that cells expressing MinC(R133A) phenocopy cells deleted for *minC*, both in morphology and Gfp-MinD oscillation.

**Fig. 2.**
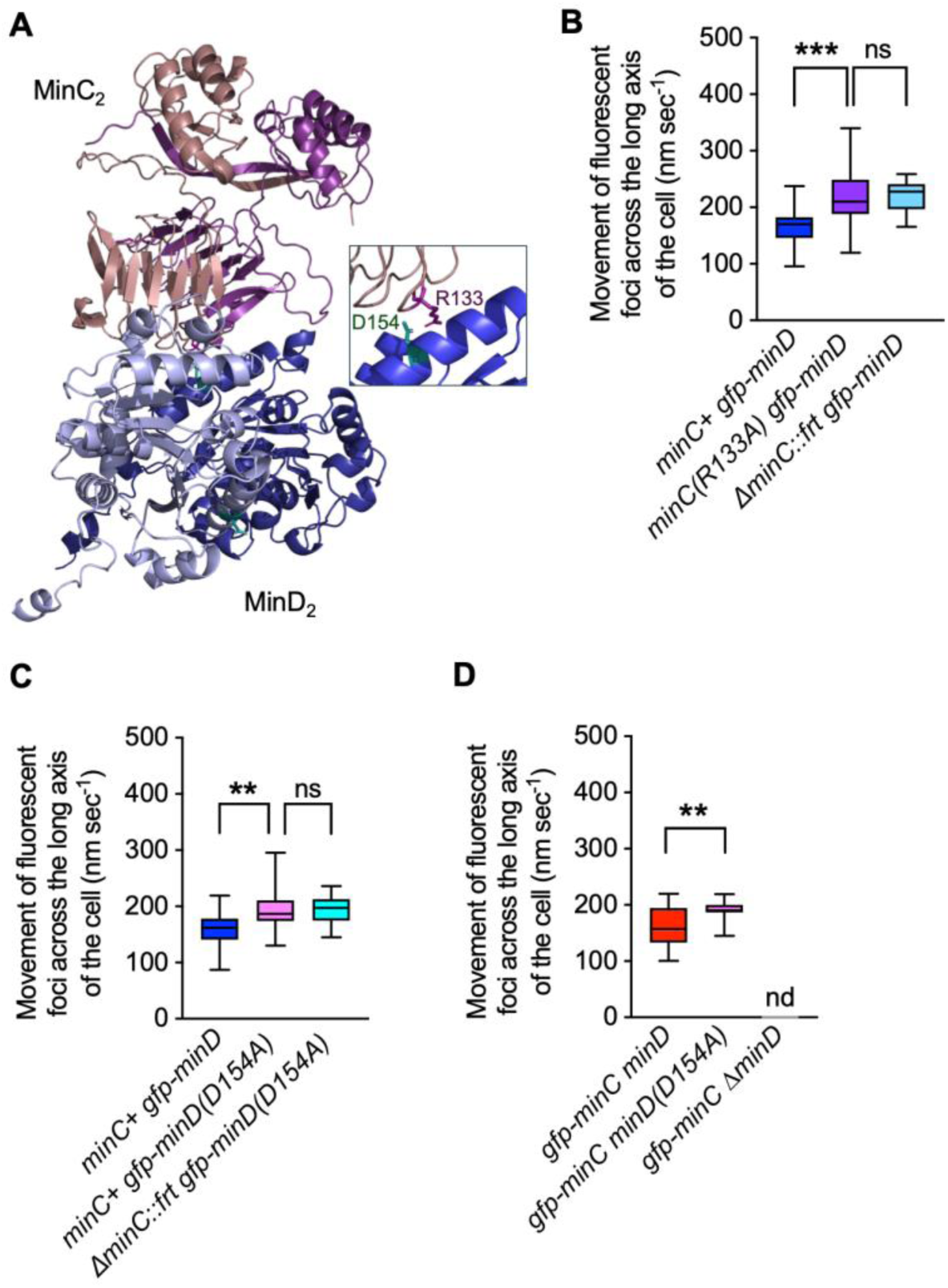
Mutations at the binding interface of MinC and MinD alter oscillation in vivo. (A) An Alphafold2 model of a MinD^2^-MinC^2^ heterodimer was constructed (MinD protomers are shown in blue and MinC protomers are shown in purple) with residues D154 of MinD and R133 of MinC highlighted in green and purple, respectively (inset) (49–52). (B) Oscillation rates of Gfp-MinD are shown in a strain expressing MinC(R133A) from the chromosome (CF095) or in a strain deleted for *minC* (CF315) (n = 20). (C) Oscillation rates of Gfp-MinD(D154A) are shown in a strain with (CF089) and without *minC* (CF313) (n = 20). (D) Oscillation rates of Gfp-MinC are shown in a strain expressing MinD(D154A) from the chromosome (CF086) or deleted for *minD* (CL0481) (n = 20). In (B-D), *p* values are as follows, ∗∗ < 0.005, *** < 0.0001, ns refers to not significant, and nd refers to no data). Error is shown as standard error.

Next, we constructed a strain containing Gfp-MinD(D154A) at the *minD* locus to disrupt the interaction on the MinD side of the interface and monitored oscillation in cells containing *minC* (CF089). We observed that Gfp-MinD(D154A) oscillates at a rate of 184.3 ± 6.0 nm sec^-1^, which is 14% faster than Gfp-MinD (Fig. 2C). To determine if deletion of *minC* from this strain increases oscillation further, we removed *minC* as before (CF313). In a strain without *minC*, we observed that Gfp-MinD(D154A) oscillates at 186.2 ± 4.2 nm sec^-1^ (Fig. 2C) (Fig. S2A). This suggests that MinD(154A) oscillation is not further modified by MinC. Cells expressing Gfp-MinD(D154A) (CF089) displayed mild filamentation and minicell formation, with the mean length of cells being 3.1 ± 0.10 µm with a minicell frequency of 6.97% (Fig. S2B) (Table 2), indicating that substitution of D154 produces a cell division defect.

We performed a reciprocal localization experiment and mutagenized the *min* locus in the chromosome to contain *gfp-minC* or, for comparison *gfp-minC(R133A)*, by site-specific recombination and then monitored oscillation in cells expressing Gfp-MinC or Gfp-MinC(R133A) [(CL0248) and (CF266), respectively]. The cells harboring *gfp-minC* (CF0248) displayed normal morphology with a mean cell length of 1.6 ± 0.09 µm and no discernable minicells. In the strain containing *gfp-minC(R133A)* (CF266), cells were mildly filamentous, with a mean length of 3.5 ± 2.34 µM, and 10.71% were minicells (Fig. S3A) (Table 2). We observed that Gfp-MinC oscillated at a rate of 165.4 ± 5.1 nm sec^-1^ (n = 20) (Fig. 2D), similar to the oscillation rate of Gfp-MinD (166 ± 7.3 nm sec^-1^) (Fig. 1A and 1C), and cells were similar in length to wild type cells (Table 2). However, Gfp-MinC(R133A) failed to oscillate and instead, cells contained cytoplasmic fluorescence (Fig. S3B) (Table 2), further supporting that Gfp-MinC(R133A) is defective for binding MinD.

Finally, we monitored Gfp-MinC oscillation in cells engineered to contain *minD(D154A)* at the *minD* locus (CF086) and in cells deleted for *minD* (CF313). In cells expressing *minD(D154A)* from the chromosome, cells displayed mild filamentation with a mean length of 2.9 ± 0.12 µM and a minicell percentage of 10.31% (Fig. S3C) (Table 2). As expected, deletion of *minD* abrogated Gfp-MinC oscillation in most cells, and Gfp-MinC fluorescence was mostly cytoplasmic (Fig. 2D) (Fig. S3A) (Table 2). Notably, 23% of cells expressing Gfp-MinC and MinD(D154A) still exhibited Gfp-MinC oscillation and, when we measured the rate of the oscillation, we observed that Gfp-MinC in cells expressing MinD(D154A) as the only copy of MinD exhibited oscillation at a rate of 188.4 ± 8.2 nm sec^-1^, nearly identical to the oscillation rate of Gfp-MinD(D154A) (Fig. 2C and 2D) (Fig. S3D) (Table 2). These results suggest that the D154A mutation does not completely block the interaction with MinC in vivo since Gfp-MinC continues to oscillate in this background. However, the strain containing Gfp-MinC and MinD(D154A) is mildly filamentous with an average cell length of 2.9 ± 0.12 µm and produces minicells (10.31%) (Fig. S3C) (Table 2), indicating that the strain is defective for normal FtsZ-ring placement. Expression of MinD variants in the Gfp-MinC background was similar among all strains tested (Fig S1E).

### MinC alters pools of membrane-associated MinD when MinE is present

When MinD oscillates between the poles of a cell, which we observe as rapid fluorescence redistribution between the poles, the behavior is produced by cycles of MinD binding to and subsequently releasing from the membrane. This is best characterized by a diffusion-capture-release mechanism (9, 29, 36). MinE is thought to regulate this process by promoting MinD ATP hydrolysis (17, 21, 29). MinE displays an oscillation that is distinct from MinD; it forms a membrane-localized ring that follows MinD across the long axis of the cell. After release, MinD cannot bind where MinE is located, so it re-binds the membrane ahead of MinE, creating a leading edge (29, 37). Moreover, the MinC and MinE binding sites on MinD partially overlap, suggesting that both cannot bind simultaneously. To understand how MinC may modify MinD membrane binding and release and slow the observed oscillation rate of Gfp-MinD from pole-to-pole in vivo, we considered two scenarios: (1) a simple competition model and (2) a MinC priming model (Fig 3A). In the simple competition model, MinC would bind MinD at the membrane effectively competing with MinE. Therefore, MinC would act to prevent release of MinD from the membrane by MinE and effectively increase membrane dwell time, slowing oscillation. Alternatively, in the MinC priming and activation model, MinC would bind MinD at the membrane and facilitate or promote recruitment of MinE and subsequent MinD release. Thus, MinC would facilitate a faster membrane binding and release cycle, create a more dynamic pool of diffuse MinD, and consequently accelerate pole-to-pole oscillation. To distinguish which of these models is plausible, we reconstituted membrane binding and release assays in vitro using purified MinC, MinD, and MinE.

**Fig. 3.**
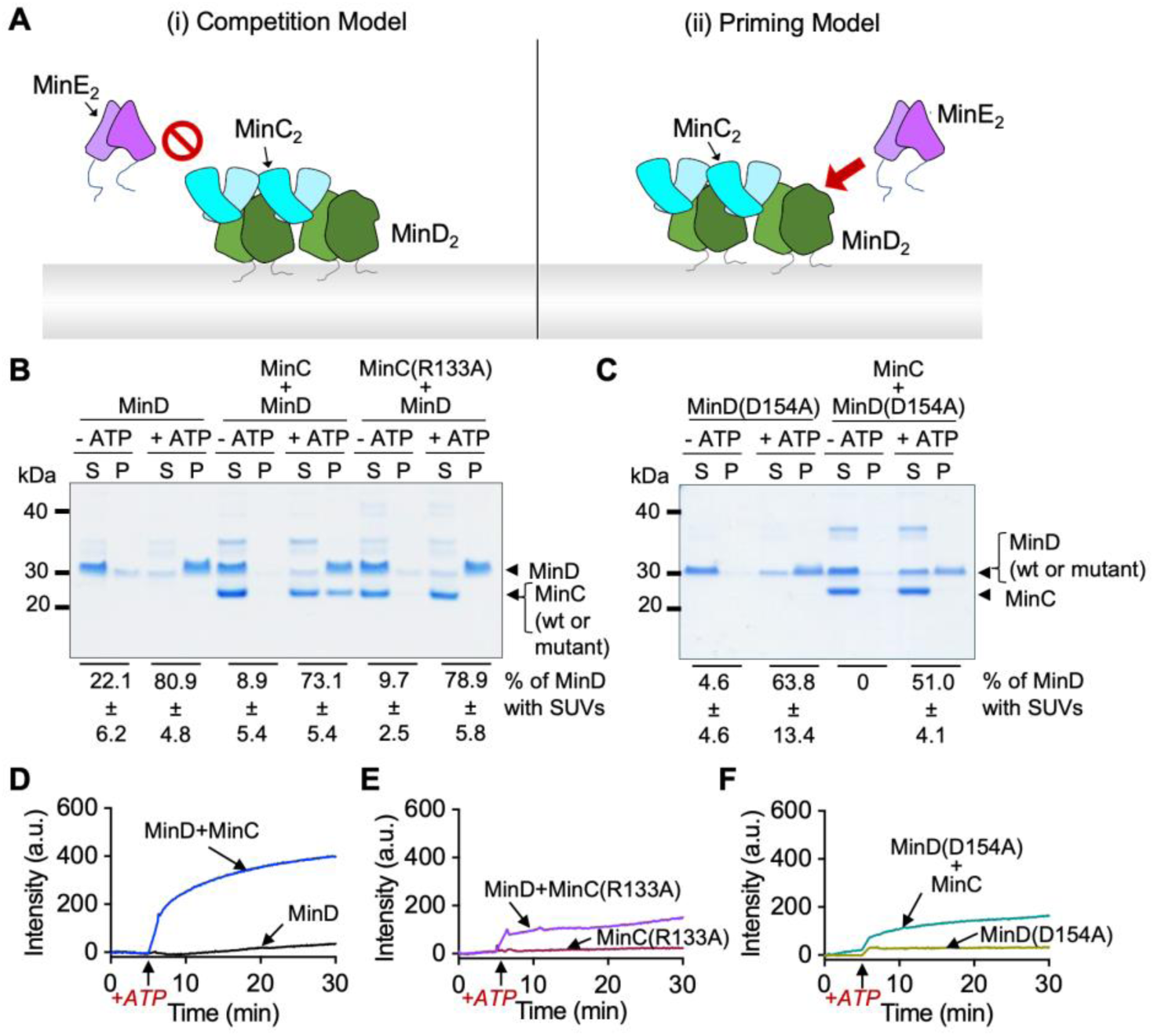
MinD and MinC binding site variants fail to interact in vitro. (A) A cartoon model illustrating two proposed mechanisms by which MinC may modulate oscillation of MinD is shown: (i) Competition Model in which there is direct competition between MinE and MinC for binding to MinD; and (ii) Priming Model in which MinC binds to and activates MinD for enhanced recruitment of MinE and subsequent release. (B) Phospholipid binding by MinD in the absence and presence of MinC was evaluated by centrifugation and SDS-PAGE of soluble (S) and SUV-associated pellet (P) fractions. MinD (4 µM) was incubated with SUVs (250 µg ml^-1^) and, where indicated, ATP (4 mM), MinC (4 µM) or MinC(R133A) (4 µM) were included in the reaction. Gel image is representative of at least three replicates. (C) MinD(D154A) (4 µM) was evaluated for binding to SUVs (250 µg ml^-1^) in reactions containing ATP and MinC as described in (B). Gel image is representative of at least three replicates. (D) To monitor assembly of copolymers with MinC, MinD (8 µM) was incubated in the absence or presence of MinC (4 µM), and light scatter was monitored for five minutes, and then ATP (8 mM) was added to the reaction. The increase in light scatter was monitored for an additional 25 minutes. (E) MinD(R133A) (4 µM) was analyzed for copolymer formation with MinD (8 µM) as described for wild type MinD in (D) by monitoring 90-degree light scatter after addition of ATP. (F) MinD(D154A) (8 µM) was analyzed for copolymer formation with MinC (4 µM) as described for wild type MinD in (D) by monitoring 90-degree light scatter after addition of ATP. Data in D through F is representative of at least three replicates. Error is shown as standard error.

First, we incubated MinD (4 µM) with small unilamellar vesicles (SUVs) in the absence and presence of MinC (4 µM) and ATP. Then, we collected soluble and SUV-associated proteins by centrifugation and analyzed both fractions by SDS-PAGE. We observed that MinD is not recruited to SUVs without ATP in centrifugation assays (Fig. 3B). However, in the presence of ATP, 80.9% of MinD associated with the phospholipid-containing SUV pellet indicating stable membrane binding (Fig. 3B) (Table 3).

**Table 3.** Phospholipid Binding of MinD and MinC Variants.

| **Reaction Conditions for**<br>**MinD wild type or variant** | **MinD localized to**<br>**PL Pellet (%)** | **MinD(D154A) localized**<br>**to PL Pellet (%)** |
| --- | --- | --- |
| PL - ATP | 22.1 ± 6.2 | 4.6 ± 4.6 |
| PL + ATP | 80.9 ± 4.8 | 63.8 ± 13.4 |
| MinC + PL - ATP | 8.9 ± 5.4 | 0 |
| MinC + PL + ATP | 73.1 ± 5.4 | 51.0 ± 4.1 |
| MinC(R133A) + PL -ATP | 9.7 ± 2.5 | n.t.<sup>1</sup> |
| MinC(R133A) + PL + ATP | 78.9 ± 5.8 | n.t.<sup>1</sup> |

<sup>1</sup> n.t. indicates not tested.

Next, we tested if the addition of MinC altered the association of MinD with SUVs and if MinD is also capable of recruiting MinC to SUVs. The addition of MinC did not significantly change the amount of MinD associating with phospholipids when ATP was present (73.1% to 80.9%), and 43.7% of MinC in the reactions was redirected to phospholipid pellets with MinD (Fig. 3B) (Table 3). We also purified and tested MinC(R133A) in phospholipid-binding assays with MinD and ATP. We observed that unlike wild type MinC, MinC(R133A) failed to localize to MinD and phospholipids when ATP was included in the reaction, consistent with Arg 133 of MinC being critical for binding to MinD (Fig. 2A) (Fig. 3B). We next tested if purified MinD(D154A) was similarly defective for recruiting MinC. MinD(D154A) exhibited reduced binding to phospholipids with ATP, with 63.8% localizing to the phospholipid pellet, and MinD(D154A) failed to recruit MinC (Fig. 3C) to the pellet (Table 3). Interestingly, Gfp-MinC oscillates in 23% of cells containing MinD(D154A), suggesting that it is capable of supporting transient interactions in vitro, although we did not detect this in reconstituted phospholipid binding assays in vitro (Fig. 2C and Fig. 3C).

MinC and MinD dimers also assemble into alternating copolymers in vitro when ATP is present, even in the absence of phospholipids, which can be monitored by measuring the 90-degree light scatter of polymer complexes or by ultracentrifugation at high speed. To monitor copolymer assembly and determine if MinC(R133A) and MinD(D154A) are defective for assembly of stable copolymers, we monitored 90-degree light scatter after addition of ATP. First, we incubated MinD (4 µM) alone, and with MinC (4 µM), measured the light scatter for 5 minutes to establish a baseline, and then added ATP (8 mM). In reactions containing MinC and MinD, we observed a rapid increase in light scattered after the addition of ATP, which plateaued after 25 minutes (Fig. 3D). However, when MinC was omitted from the reactions, no increase was detected after ATP was added. Next, we repeated the assay but substituted MinC(R133A) for wild type MinC. We detected a small increase in light scatter in reactions containing MinD, MinC(R133A) and ATP (Fig. 3E), but the amplitude was 63% less than what was observed for wild type MinC. Finally, MinD(D154A) also exhibited a small increase in light scatter in reactions containing MinC after the addition of ATP, but was 64% less than what was observed with wild type MinD (Fig. 3F). Together, these results show that wild type MinC and MinD assemble into stable copolymers with ATP, but MinC(R133A) and MinD(D154A) are largely impaired for ATP-dependent copolymer formation. To confirm these results, we also performed ultracentrifugation assays to collect and analyze MinC-MinD copolymers. We again observed that both MinC(R133A) and MinD(D154A) are defective for assembly of stable copolymers, compared with their wild type counterparts, after addition of ATP (Fig. S4) (Table S3). Together, these results further confirm that Arg 133 of MinC and Asp 154 of MinD are important residues in the MinC-MinD interface and support stable complex formation and copolymers.

To reconstitute the membrane release step of MinD in vitro, we performed a staged, sequential recruitment assay in which we added MinE to MinD or to MinC-MinD that was pre-loaded onto SUVs. First, we preassembled SUVs (250 µg ml^-1^) with MinD (4 µM) and ATP (4 mM), in the presence and absence of MinC (4 µM) for five minutes, and then MinE (16 µM) was added to the reaction (Fig. 4A). Phospholipid-bound proteins were collected by centrifugation (21,000 x *g*) for 15 minutes, and then supernatants and SUV pellets were analyzed by SDS-PAGE. In the absence of MinE, 82.9 ± 5.3% of MinD fractionated with SUVs when ATP was present; however, in the presence of MinE, the majority of MinD was released to the supernatant and only 22.7 ± 0.6% of MinD remained in association with SUV’s (Fig. 4B) (Table 4). When MinC was included in the reaction with MinD, we saw a further reduction of SUV-bound MinD, to 13.8 ± 4.1% of MinD associated with phospholipids (Fig. 4B). This suggests that MinC fails to prevent, but rather may enhance, MinE-stimulated release of MinD from phospholipids.

**Fig. 4.**
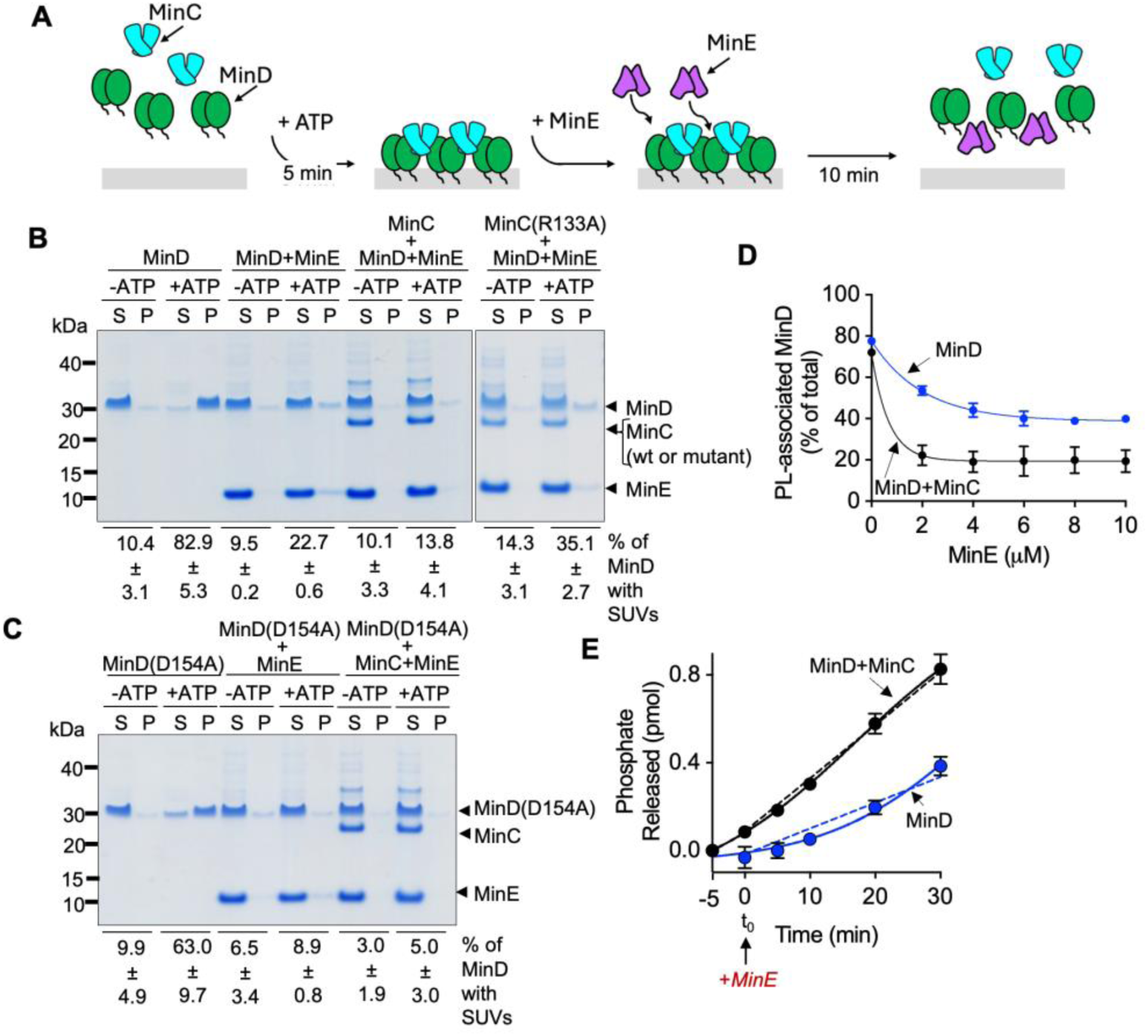
Staged reconstitution assays suggest MinC-directed activation of MinD. (A) Cartoon schematic of a staged ATP hydrolysis assay is shown where MinD, MinC and ATP are preloaded onto SUVs for 5 minutes, and then MinE is added to detect MinD release from the membrane and ATP hydrolysis activity. (B) Phospholipid binding by MinD and MinC with ATP; then MinD release by MinE was evaluated by centrifugation and SDS-PAGE of soluble (S) and SUV-associated pellet (P) fractions. MinD (4 µM) was incubated with SUVs (250 µg ml^-1^) and, where indicated, ATP (4 mM), MinC (4 µM) or MinC(R133A) (4 µM), and MinE (4 µM) were included in the reaction. Gel image is representative of at least three replicates. (C) MinD(D154A) (4 µM) was evaluated for binding to SUVs (250 µg ml^-1^) in reactions containing ATP and MinC as described in (B). Gel image is representative of at least three replicates. (D) Plot of the percent of total MinD associated with SUVs after addition of increasing MinE (0 µM to 10 µM) preassembled with (black line) and without (blue line) MinC. (E) Staged ATP hydrolysis assays where MinD (8 µM) pre-incubated with (black line) or without MinC (blue line) (4 µM) is preincubated with ATP (4 mM) and phospholipids (250 µg ml^-1^) for 5 minutes at 30°C, then MinE (16 µM) is added, and phosphate release is monitored for 30 minutes. Dashed lines indicate a linear model fit of activity for comparison. Data in B through E is representative of at least three replicates. Error is shown as standard error.

**Table 4.** Phospholipid Bound MinD and MinC Variants after Release by MinE.

| Reaction Conditions for MinD wild type or variant | MinD remaining with PL Pellet (%) | MinD(D154A) remaining with PL Pellet (%) |
| --- | --- | --- |
| PL - ATP | 10.4 ± 3.1 | 9.9 ± 4.9 |
| PL + ATP | 82.9 ± 5.3 | 63.0 ± 9.7 |
| PL - ATP + MinE | 9.5 ± 0.2 | 6.5 ± 3.4 |
| PL + ATP + MinE | 22.7 ± 0.6 | 8.9 ± 0.8 |
| PL - ATP + MinC + MinE | 10.1 ± 3.3 | 3.0 ± 1.9 |
| PL + ATP + MinC + MinE | 13.8 ± 4.1 | 5.0 ± 3.0 |
| PL - ATP + MinC(R133A) + MinE | 14.3 ± 3.1 | n.t.<sup>1</sup> |
| PL + ATP + MinC(R133A) + MinE | 35.1 ± 2.7 | n.t.<sup>1</sup> |

<sup>1</sup> n.t. indicates not tested.

To determine if the enhanced MinD release by MinE is attributable to MinC and occurs via the MinC-MinD interaction, we replaced MinC with MinC(R133A) in staged release assays with MinD, MinE, and ATP. After centrifugation, we observed that 35.1 ± 2.7% of MinD remained in association with phospholipids in reactions containing MinE and MinC(R133A), with no added benefit to release by MinC(R133A) (Fig. 4B) (Table 4). While the fractionation differences that we observed with and without MinC are subtle, they are consistent with a model in which wild type MinC enhances release of MinD from phospholipids by MinE, but MinC(R133A) does not. Finally, we also tested if MinD(D154A) was capable of MinE-stimulated release, since the MinC and MinE binding sites partially overlap on MinD. We observed that MinD(D154A) is partially defective for binding to phospholipids with ATP, 63.0 ± 9.7% (Fig. 4C) (Table 4). Furthermore, MinE addition promotes efficient release of MinD(D154A), and the presence of MinC does not alter this result (Fig. 4C).

To determine if MinC enhances release of MinD from phospholipids across a range of MinE concentrations, we titrated the amount of MinE (0 to 10 µM) in phospholipid release assays with MinD. We observed that MinD dissociated from phospholipids at a lower MinE concentration when MinC was present in the reaction than when MinC was omitted, with the effective concentration of half-maximal response (EC_50_) values of 0.5 µM and 4.0 µM of MinE, respectively. The lower EC_50_ of MinE in reactions containing MinC suggests that MinE is more effective at releasing MinD when MinC is present (Fig 4D). Finally, we predicted that MinC(R133A) would be incapable of enhancing MinE-stimulated release of MinD from the membrane, therefore we repeated the experiment but replaced wild type MinC with MinC(R133A). As expected, the presence of MinC(R133A) was unable to lower the EC_50_ of MinE to the extent of wild type MinC (Fig S3E) [EC_50_ with MinC(R133A) is 2.9 µM].

Stimulation of MinE function by MinC is consistent with a model in which MinC primes or activates membrane-associated MinD prior to MinE binding and inconsistent with the simple direct competition model (Fig. 3A). If MinC activates or primes MinD for MinE, then this likely occurs by stabilizing a change in the conformation of MinD on the membrane. If this is the case, and MinD-dependent release is a consequence of ATP hydrolysis, then MinC should stimulate MinE-activated ATP hydrolysis in enzyme activity assays in vitro. To test this, we performed a staged ATP hydrolysis assay where MinD, MinC, and ATP are pre-incubated together for five minutes before MinE is added, and total phosphate released is monitored over a 30 minute period. We observed that in the absence of MinC, there was a noticeable lag of approximately 15 minutes prior to the start of linear ATP hydrolysis activity. A lag in MinD ATP hydrolysis activity in vitro was previously reported by others, including the Lutkenhaus and RayChaudhuri groups (21, 38). Interestingly, this lag was not present when MinC and MinD were co-incubated together prior to the reaction (Fig. 4E) (21, 38). This is consistent with the hypothesis that MinC modifies the membrane binding and MinE-stimulated release of MinD. Accordingly, pre-incubating MinD with MinC(R133A) does not overcome the lag in hydrolysis (Fig. S3F).

### Reduced membrane binding by MinD perturbs oscillation and association with MinC

If MinC modifies the dynamic flux of MinD on and off the membrane, then it should be possible to induce changes in pole-to-pole oscillation through modifying the strength of the membrane interaction, which would impact the dwell time of MinD on the membrane without modifying the MinC interface. To test this, we calculated the membrane binding potential of the MinD MTS, an amphipathic helix containing MinD residues 253 through 270 (RFIEEE KGFLKRLFGG), using a trained model to calculate the free energy associated with membrane binding (PMIpred) (39). Based on this model, the MTS of MinD is predicted to strongly bind to a membrane (Fig. 5A). To identify an MTS variant with a weaker predicted membrane interaction, we calculated the membrane binding potentials for several MTS variants containing amino acid substitutions, including MinD(KR265AA), MinD(KR265EE), MinD(FL263GG), and a truncated variant MinD(Δ8). Several variants were predicted to lose the interaction, including MinD(Δ8) and MinD(FL263GG) (Fig. 5A). Consistent with this prediction, removal of the MTS has previously been reported to abolish membrane localization (40, 41). MinD(KR265AA) and MinD(KR265EE) were predicted to have weaker membrane interactions, with MinD(KR265EE) potentially losing binding altogether (Fig. 5A). To test if these variants were functional, we constructed chimeric Gfp fusion proteins and introduced them into the chromosome at the *min* locus. All four variants, even the most conserved mutant MinD(KR265EE) induced filamentation, with minicells present, indicating a cell division defect (Fig. 5B) (Fig. S6A) (Fig. S7A) (Table 2). When we monitored oscillation of Gfp-MinD chimeras, only Gfp-MinD(KR265AA) exhibited measurable oscillation (Fig. 5C and 5D). Gfp-MinD(KR265AA) oscillated at a 14% faster rate than Gfp-MinD (185.7 ± 9 nm sec^-1^) (Table 2) (Fig 5C and Fig. 5E). Of the other variants tested, Gfp-MinD(KR265EE), Gfp-MinD(FL263GG), and Gfp-MinD(Δ8), all failed to oscillate. All variants were expressed to similar levels as Gfp-MinD (Fig S7D). Next, we performed the reciprocal experiment with the strain containing chromosomally located *gfp-minC* and *minD* variants in the chromosome. Again, expression of the variants produced filamentous cells and minicells, suggestive of a cell division defect (Table 2). Gfp-MinC failed to oscillate in strains containing MinD(FL263GG), MinD(KR265EE), or MinD(Δ8) (Table 2) (Fig S7B and S7C). However, Gfp-MinC, in the strain containing MinD(KR265AA), oscillated at an accelerated rate, similar to the strain expressing Gfp-MinD(KR265AA), 191.8 ± 11.1 nm sec^-1^ and 185.7 ± 9.0 nm sec^-1^, respectively (Fig 5D) (Fig 5F) and faster than wild type MinD.

**Fig. 5.**
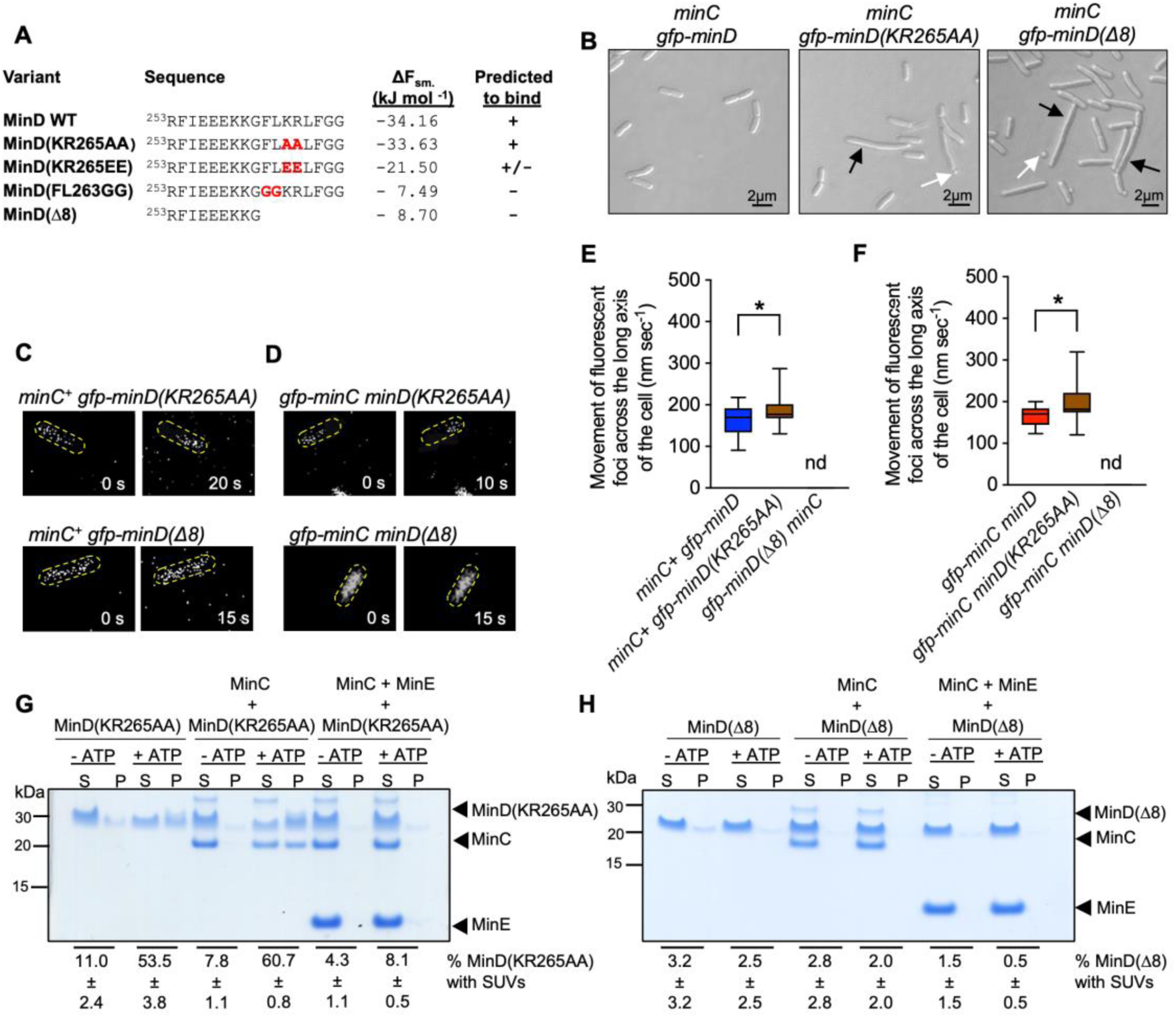
Changes to the MinD MTS alter oscillation of Gfp-MinD. (A) MinD MTS variants are shown with the calculated membrane binding free energy (kJ mol^-1^) (ΔF_sm). Predicted binding is denoted next to the MinD variant shown by either ‘+’ for binding, ‘+/-’ for intermediate, or ‘–’ for no binding for a vesicle with a radius of 50 nm. (B) DIC microscopy was performed to compare morphology of strains containing *gfp-minD, gfp-minD(KR265AA), and gfp-minD(Δ8).* Black arrows denote filamentous cells, and white arrows denote minicells. Scale bar is 2 µm. (C) Confocal fluorescence microscopy images (time lapse) show oscillating polar fluorescence by Gfp-MinD(KR265AA) in live cells (CF097) and non-oscillating diffuse cytoplasmic fluorescence of Gfp-MinD(Δ8) (CF256). (D) Time lapse fluorescence images as in (C) in strains expressing Gfp-MinC and MinD(KR265AA) (CF071) or MinD(Δ8) (CF070). (E) Oscillation rate of Gfp-MinD(KR265AA) was measured and compared to Gfp-MinD. The strain expressing Gfp-MinD(Δ8) did not exhibit Min oscillation (F) Oscillation rate of Gfp-MinC in a strain containing MinD(KR265AA) was monitored and compared to a strain containing wild type MinD. (G) Phospholipid binding assay of MinD(KR265AA) (4 µM) with and without ATP (4 mM) and MinC (4 µM). Fraction bound and released by MinE (16 µM) was evaluated by centrifugation and SDS-PAGE of soluble (S) and phospholipid-associated (P) fractions. Gel is a representative image of at least three replicates. (H) Phospholipid binding assay of MinD(Δ8) (4 µM) with and without ATP (4 mM) and MinC (4 µM). Fraction bound and released by MinE (16 µM) was evaluated by centrifugation and SDS-PAGE of soluble (S) and phospholipid-associated (P) fractions. Data in E through H are representative of at least three replicates; *p* values are as follows, ∗ ≤ 0.01, and ‘nd’ refers to no data. Error is shown as standard error.

To evaluate binding of MinD(KR265AA) to phospholipids, we purified the protein and performed phospholipid recruitment assays in vitro. We observed that MinD(KR265AA) binds to SUVs when ATP is present with 53.5 ± 3.8% of the protein associated with SUVs. However, the proportion bound to SUVs is significantly lower than wild type MinD (Table 4), suggesting that MinD(KR265AA) has a minor phospholipid binding defect (Fig 5G), as predicted by the model (Fig 5A). This suggests that MinD(KR265AA) would likely have a higher fraction of protein in the cytoplasm capable of diffusion. The other three variants were mostly defective for binding to SUVs (Table 5) (Fig. S6E and S6F). We next added MinC to the staged reactions to determine if MinD(KR265AA) recruited MinC to SUVs. We observed that MinD(KR265AA) efficiently recruited MinC to SUVs (Fig. 5G). The other variants were defective for recruiting MinC to SUVs (Fig. 5H) (Fig. S6E and S6F). Interestingly, all variants were partially defective for ATP hydrolysis with MinE and SUVs (Fig S6B). We next tested if MinD(KR265AA) forms copolymers with MinC and ATP. However, we observed that all variants tested, including MinD(KR265AA), were unable to form copolymers in vitro (Fig. S5C-F) (Fig. S6C-F). This suggests that association into higher order structures is, at least in part, facilitated by the MTS of MinD.

**Table 5.** Phospholipid Binding of MinD and MinC Variants and Release by MinE.

| Reaction Conditions for<br>MinD wild type or variant | MinD(KR265AA)<br>localized to PL<br>Pellet (%) | MinD(KR265EE)<br>localized to PL<br>Pellet (%) | MinD(FL263GG)<br>localized to PL<br>Pellet (%) | MinD(Δ8)<br>localized to PL<br>Pellet (%) |
| --- | --- | --- | --- | --- |
| PL - ATP | 11.0 ± 2.4 | 4.9 ± 1.7 | 2.7 ± 0.8 | 3.2 ± 3.2 |
| PL + ATP | 53.5 ± 3.8 | 17.6 ± 9.6 | 4.3 ± 0.5 | 2.5 ± 2.5 |
| PL - ATP + MinE | 7.8 ± 1.1 | 3.8 ± 0.9 | 5.3 ± 3.2 | 2.8 ± 2.8 |
| PL + ATP + MinE | 60.7 ± 0.8 | 19.0 ± 2.0 | 1.7 ± 0.6 | 2.0 ± 2.0 |
| PL - ATP + MinC + MinE | 4.3 ± 1.1 | 1.6 ± 0.9 | 3.1 ± 0.7 | 1.5 ± 1.5 |
| PL + ATP + MinC + MinE | 8.1 ± 0.5 | 3.6 ± 2.3 | 1.9 ± 1.5 | 0.5 ± 0.5 |

## Discussion

### MinC modulates oscillation in vivo by priming MinD

MinD and MinE are sufficient to establish the patterning that underpins oscillation; consequently, the role of MinC in this context has largely been attributed as cargo for the oscillation. To further understand the impact and functional role of MinC during Min system oscillation in live cells, we deleted *minC* from an *E. coli* strain engineered to contain *gfp-minD* at the *min* locus and monitored Min oscillation in live dividing cells. We observed a faster Min oscillation rate in the *minC* deletion strain than in the strain containing *gfp-minD* and *minC*. This impact of MinC on the oscillation was further validated by analyzing Min oscillation in strains expressing a variant of MinD or MinC, MinD(D154A) and MinC(R133A), respectively. Both variants were previously reported to be impaired for the interaction between MinC and MinD but not for cell viability (7). In a strain expressing *gfp-minD(D154)*, or in a strain containing *minC(R133A)* and *gfp-minD,* both strains displayed faster oscillation rates compared to a wild type *minC* strain with *gfp-minD*. These results suggest that the interaction between MinC and MinD modifies oscillation in dividing cells. This is also consistent with a previous report that showed that expression of several variants of MinC conferred changes to the overall oscillation rate (8).

It is well established that the oscillation of MinD is regulated by membrane binding and release by MinE (17–19, 37, 38). Thus, we tested the ability of MinC-MinD interface mutants, including MinD(D154A) and MinC(R133A), to associate with and release from SUVs in vitro. Using purified proteins, we observed that MinD(D154A) had a decreased ability to bind to phospholipids compared to wild type and was unable to recruit wild type MinC. This result was surprising because residue D154 is not located near the MTS of MinD and implies that disruption to the opposite face could regulate MinD membrane binding. With MinE, which promotes MinD ATP hydrolysis and release, we observed a significantly lower EC_50_ for MinE in phospholipid release assays by MinD when MinC was present than when MinC was omitted. Staged ATP hydrolysis assays further suggested that pre-incubation of MinC with MinD activates MinD for MinE binding, ATP hydrolysis, and release, thus overcoming a perceived lag in reconstituted activity assays, which had previously been reported in the literature (21, 38).

### MinC alters the diffusive and membrane-associated ratios of MinD

What is perceived as pole-to-pole oscillation is best described by diffusion-capture-release of MinD on the membrane (20, 21, 36–38, 40, 42, 43). Thus, there are two distinct populations of MinD, a membrane bound, ATP-associated population, as well as a cytoplasmic pool, which is subject to diffusion. We hypothesize that through altering the efficiency of the committed MinD release step, MinC exerts control over the ratio of MinD present in the cytoplasmic and membrane pools. Strains deleted for *minC* exhibit a faster oscillation rate, which may be explained by a larger diffusive fraction of MinD as MinC accelerates the highly dynamic diffusion-capture-release cycle. Passive diffusion is approximately one hundred times faster than Min oscillation (41, 44).

We also identified a MinD MTS variant with a reduced capacity for membrane association, and expression of this variant supported oscillation, but the oscillation rate was faster than wild type as predicted by our model. This result is consistent with oscillation rates and timing subject to the dynamic flux of cytoplasmic and membrane pools of MinD. A previous report described Min patterning as being a toggle switch controlled by the ratio of MinD-to-MinE on the membrane and characterized by dynamic depletion or MinD from the cytoplasm (45). Our work extends this proposed mechanism by demonstrating that MinC regulates the diffusive pool of MinD by modifying the ratio of MinD in the membrane bound state and the cytoplasmic pool, which is in dynamic flux and subject to displacement by MinE. Therefore, we show that in addition to destabilizing FtsZ polymers, MinC functions to prime MinD on the membrane for more efficient release by MinE, thereby exerting a regulatory effect on oscillation dynamics. We predict that this occurs through MinC inducing or stabilizing a conformational change in MinD that favors MinE binding, ATP hydrolysis, and release. MinC would thus act to adjust the stoichiometric balance between the pools of MinD, cytoplasmic and membrane bound, and commit MinD to MinE activation. This mechanism also partly explains why changes to the stoichiometric ratios of MinC, MinD and MinE lead to a breakdown in the oscillation. A more intriguing question at this time is why oscillation is tunable through MinC overall concentration and direct interactions. Is there a survival advantage provided to bacteria that exhibit adaptable oscillation systems? Evolution has preserved the oscillation across a wide range of organisms-although one could envision simpler gradient systems to delineate middles and poles, if that were the sole purpose.

## Material and Methods

### Bacterial strains, plasmids, and growth conditions

*E. coli* strains, as shown in Table 1, were grown in LB (Lennox) broth, or on LB agar (Lennox) as indicated, supplemented with ampicillin (100 μg ml^-1^), chloramphenicol (35 μg ml^-1^) and kanamycin (50 μg ml^-1^), where appropriate. Gene deletions and insertions were be constructed by λ-Red recombination as previously described (8, 46). Site-directed mutagenesis of plasmids containing *minC*, *minD*, and *minE* was performed using the QuikChange II XL mutagenesis system (Agilent) and confirmed by Sanger sequencing (Table S1 and Table S2). Densities of cultures was monitored by OD_600_. Cells encoding *gfp-minC* were grown in the presence of 100 mM L-arabinose to induce expression. Plasmid pET*-minD* was mutagenized to construct the following for purification: pET-*minD(FL263GG)*, pET-*minD(KR265AA)*, pET-*minD(KR265EE)*, pET-*minD(Δ8)*, and pET-*minD(D154A)*. Plasmid pET-*minC* was mutagenized to construct pET-*minC(R133A)* for purification. Plasmid pBAD33-*gfp-minD* was mutagenized to construct the following: pBAD33-*gfp-minD(FL263GG)*, pBAD33-*gfp-minD(KR265AA)*, pBAD33-*gfp-minD(KR265EE)*, pBAD33-*gfp-minD(Δ8)*, and pBAD33-*gfp-minD(D154A).* The plasmid pBAD33-*gfp-minC* was mutagenized to construct pBAD33-*gfp-minC(R133A).* Plasmids (pBAD) were used to amplify genes prior to site-specific insertion in the chromosome. All strains were confirmed by sequencing.

### Fluorescence Microscopy

Wild type, overexpression, and deletion strains containing *gfp–minC and gfp–minD* genes (Table 1), where indicated, were grown and imaged by differential interference contrast (DIC) microscopy using a Nomarski prism with a Zeiss AxioCam HRc high-resolution camera. Images were prepared using Adobe Photoshop 2024, and cell lengths were measured using NIH ImageJ software. To observe oscillation in live cells, cells were grown to log phase (OD_600_ of 0.5) at 30°C and applied to agarose gel pads (5% w/vol) containing M9 minimal medium supplemented with 0.2% glucose. Live cells were visualized by confocal fluorescence microscopy with a Zeiss AxioImager M2 system and LSM 700 confocal module with excitation and emission wavelengths set to 488 and 525. Oscillation rate was measured by identifying the fluorescence focal center of a single cell using NIH ImageJ and tracking foci movement across the longitudinal cell axis (nm sec^−1^) in a time series of live cell images, collected in 5 second intervals for 30 frames. This was performed for at least 20 single cells per strain visualized. The peak polar fluorescence was determined using NIH ImageJ software at one pole and tracked to the opposite pole then plotted as a function of distance traveled with time to calculate rate (nm sec^-1^). Kymographs were constructed by plotting total fluorescence across the longitudinal axis a single cell as a time series from live cell images taken every 5 seconds for at least 12 frames.

### Protein extraction and Immunoblotting

Wild type and mutant strains were grown to log phase in glucose minimal medium as indicated at 30°C. Culture samples (1 ml) were collected, and cells were pelleted by centrifugation at 6,000 x *g*. Protein was extracted by the addition of bacterial protein extraction reagent (B-PER) (Thermo Scientific) and 20 μg ml^-1^ of lysozyme, and protein concentration was determined by the bicinchoninic acid (BCA) assay. Protein samples were analyzed by reducing SDS-PAGE and transferred to a nitrocellulose. Membranes were washed with Tris-buffered saline (pH 7.6) and Tween-20 (0.05%), blocked for 2 h with 5% (w/v) bovine serum albumin, probed with anti-MinD rabbit polyclonal serum (1:5000 dilution) and anti-rabbit IgG coupled with horseradish peroxidase (1:20,000 dilution). MinD was visualized using SuperSignal West Pico Plus chemiluminescent substrate (ThermoFisher).

### Expression and purification of proteins

MinC, MinD, and MinE proteins were overexpressed from *E. coli* BL21Δ*clpP::cat* harboring pET28a-*minC*, pET28a-*minD,* pET28a-*minE* respectively. Proteins were purified as previously reported (27). Briefly, MinC, MinD, and MinE were each expressed from plasmids by adding isopropyl β-d-1-thiogalactopyranoside (IPTG) (0.5 mM). Cells were collected and lysed by French press. Briefly, soluble cell extracts were added to a Q Sepharose column and proteins were eluted with a linear gradient of KCl (50 mM to 600 mM). Fractions containing the proteins of interest were fractionated on an S100 column. MinC(R133A), MinD(D154A), MinD(FL263GG), MinD(KR265AA), MinD(KR265EE), and MinD(Δ8) mutant proteins were constructed via site-directed mutagenesis of *minC, minD* or *minE* expression plasmids. All protein concentrations are reported as MinC, MinD, and MinE dimers.

### Phospholipid binding and release

Phospholipid recruitment assays were performed using MinD (4 μM), MinD (D154A) (4 μM), MinD (KR265AA) (4 μM), or MinD (FL265GG) (4 μM), in assembly buffer containing 50 mM 2-(*N*-morpholino)ethanesulfonic acid (MES) pH 6.5, 100 mM KCl, 10 mM MgCl_2_ (25 μl) supplemented with SUVs (250 μg ml^-1^). Where indicated, MinC (0 μM to 6 µM) or MinC (R133A) (4 µM), MinE (16 µM), and ATP (4 mM) were included. Reactions were incubated for 10 minutes at 30 °C, following the addition of ATP, where indicated. SUV-associated proteins were collected by centrifugation at 21,000 x *g* at 23 °C for 15 minutes. Supernatant and pellets were resuspended in equivalent volumes of buffer and 1X lithium dodecyl sulfate (LDS) sample buffer and analyzed by SDS-PAGE. After staining, densitometry was performed to quantitate band intensity using NIH Image J. SUVs from *E. coli* extracts (Avanti Polar lipids) were prepared as previously reported (47). Phospholipid release was monitored similarly, but reactions were supplemented with MinE, where indicated (0 μM to 10 µM), and incubated for 10 minutes at 30 °C.

### Light Scatter Assays to Monitor for MinC and MinD Copolymer

Formation of MinC-MinD copolymers was monitored at 23 °C by 90° light scatter with excitation and emission wavelengths set to 450 nm. Reactions containing MinC (4 µM) or MinC(R133A) (4 µM) was added to reactions containing MinD (8 µM), MinD (D154A) (8 µM), MinD (KR265AA) (8 µM), MinD (KR265EE) (8 µM), or MinD (FL263GG) (8 µM) in copolymer assembly buffer (20 mM MES, pH 6.5, 100 mM KCl, 10 mM MgCl_2_). ATP (8 mM) was added to reactions after 5 minutes and light scattered was monitored for an additional 30 minutes.

### MinD and MinC Copolymer Formation Ultracentrifugation Assay

MinC (4 µM) or MinC(R133A) (4 µM) was added to reactions containing MinD (8 µM) or MinD (D154A)(8 µM) in assembly buffer containing (20 mM MES, pH 6.5, 100 mM KCl, 10 mM MgCl_2_) as indicated. Reactions were incubated for 15 min at 23 °C following the addition of ATP (4 mM), then centrifuged at 129,000 x *g* for 30 minutes at 23 °C. Supernatants and pellets were resuspended in 1X LDS sample buffer, analyzed by SDS-PAGE. Band intensity and quantified using NIH ImageJ.

### ATP hydrolysis assays

ATP hydrolysis was measured by monitoring the amount of inorganic phosphate released after 10 min in reactions containing Tris buffer (20 mM, pH 7.5), KCl (50 mM), MgCl_2_ (10 mM), ATP (4 mM), MinD (8 μM), MinD(D154A) (8 μM), MinD(KR265AA) (8 μM), MinD(KR265EE) (8 μM), MinD (FL263GG) (8 μM), SUVs (250 μg ml^-1^), and MinE (16 μM), where indicated. Staged ATP hydrolysis assays were performed with MinD (8 μM), MinE (16 μM), and SUVs (250 μg ml^-1^), with or without MinC or MinC (R133A) (4 µM). Staged assays were conducted by preincubating MinD (8 µM) with MinC (4 µM) and ATP (4 mM) for 5 minutes before addition of MinE (16 µM). Total inorganic phosphate was measured at 5,10, 20, and 30 minutes. Phosphate released in reactions was detected using Biomol green (Enzo Life Sciences) and compared to a phosphate standard curve.

## Acknowledgements

We thank Josiah Morrison, Ben Piraino, Ellen Madden, Kate Remy, and Jake Grecco for helpful discussions and critical reading of the article and Janet Atoyan for microscopy and sequencing technical assistance. Research reported in this publication was supported by the National Institute of General Medical Sciences of the National Institutes of Health under Award Number R01GM158901 to J. Camberg. The content is solely the responsibility of the authors and does not necessarily represent the official views of the National Institutes of Health.

## Data availability

All data pertinent to this work are contained within this manuscript or available upon request. For requests, please contact Jodi Camberg at the University of Rhode Island.

## Supplementary Materials

Supplementary Tables S1, S2, and S3 and Supplementary Figures S1, S2, S3, S4, S5, S6, and S7 are provided as a separate file.

