## Supplementary Materials for "MinC influences the Min oscillation period in *Escherichia coli* by priming MinD on the membrane"

Submitted to BioRXIV October 5, 2026

### Supplementary Tables

#### Table S1. Plasmid List

| Plasmid | Antibiotic selection marker | Source, reference, or construction |
|---|---|---|
| pKD3 | *cat* | (46) |
| pKD46 | *amp* | (46) |
| pKD267 | *kan (kan-$P_{rha}$-parE)* | (8) |
| pCP20 | *amp* | (53) |
| pBad33 | *cat* | (53) |
| pBad33-*gfp-minD* | *cat* | (28) |
| pBad33-*gfp-minD(FL263GG)* | *cat* | This study |
| pBad33-*gfp-minD(KR265AA)* | *cat* | This study |
| pBad33-*gfp-minD(KR265EE)* | *cat* | This study |
| pBad33-*gfp-minD(Δ8)* | *cat* | This study |
| pBad33-*gfp-minD(D154A)* | *cat* | This study |
| pBad33- *$P_{BAD}$-gfp-minC* | *cat* | (8) |
| pBad33- *$P_{BAD}$-gfp-minC(R133A)* | *cat* | This study |
| pET28a | *kan* | MilliporeSigma, US |
| pET28a-*minC* | *kan* | (32) |
| pET28a-*minC(R133A)* | *kan* | This study |
| pET28a-*minD* | *kan* | (32) |
| pET28a-*minD(FL263GG)* | *kan* | This study |
| pET28a-*minD(KR265AA)* | *kan* | This study |
| pET28a-*minD(KR265EE)* | *kan* | This study |
| pET28a-*minD(Δ8)* | *kan* | This study |
| pET28a-*minD(D154A)* | *kan* | This study |
| pET28a-*minE* | *kan* | (32) |

### Table S2. Primer List

| Primer Name | Primer Description | Sequence |
|---|---|---|
| MinCKOGDF | Deleting *minC* from *gfp-minD* background | 5' AACATCATCG CGCGCTGGCG ATGATTAATA GCT AATTGAGTAAGGCCAGGTCCCGCTCAGAAGAACTCGT 3' |
| MinCKOGDR | Deleting *minC* from *gfp-minD* background | 5' AGTGAAAAGT TCTTCTCCTT TACTCATAGA AAT TCCTTGT TAAAAAGGGA GGATCCAATCAGCCCTTGAG 3' |
| GfpMinDKIF | To insert *gfp-minD* into native locus | 5' ACCGTT AAATTGATCC CTTTTTAACA AGG AATTTCT ATG AGT AAA GGA GAA GAA CTT TTC 3' |
| GfpMinDKIR | To insert *gfp-minD* into native locus | 5'TTTTCTTCCGCGAGAGAAAGAAATCG AGTAATGCCATAAC TTATCCTCCGAACAAGCG 3' |
| GfpMinDKIMTSR | To insert *gfp-minD* MTS variants into native locus | 5' TTTTCTTCCGCGAGAGAAAGAAAT CGAGTAATGCCATAAC TTATCCTCCGAACAA 3' |
| MinCKOCatF | To delete *minC* FRT sites | 5' AACATCATCG CGCGCTGGCG ATGATTAATA GCTAAT TGAG TAAGGCCAGG GTGTAGGCTGGAGCTGCTTC 3' |
| MinCKOCatR | To delete *minC* FRT sites | 5'AGTGAAAAGTTCTTCTCCTTTACTCATAGAAATTCCTT GTTAAAAAGGGA CATATGAATATCCTCCTTAGTTCC 3' |
| pbadGfpMinCKIF | To insert *gfp-minC* at the native locus | 5' GCTGGCGATGATTAATAGCTAATTGAGTAAGGC CAGGATGTACCCGTTTTTTTGGGC 3' |
| pbadGfpMinCKIR | To insert *gfp-minC* at the native locus | 5' ATTCCTTGTTAAAAAGGGATCAATTTAACGGTTGAA CGGTCAAAGCGTTTTCGACTAAC 3' |
| MinDFL263GGF | To mutagenize residues 263 and 264 from FL to GG | 5' GAA GAG AAG AAA GGC GGC GGC AAA CGC TTG TTC GGA GGA 3' |
| MinDFL263GGR | To mutagenize residues 263 and 264 from FL to GG | 5' TCC TCC GAA CAA GCG TTT GCC GCC GCC TTT CTT CTC TTC 3' |
| MinDKR265AAF | To mutagenize residues 265 and 266 of MinD from KR to AA | 5' GAG AAG AAA GGC TTC CTC GCG GCG TTG GGA GGA TAA 3' |
| MinDKR265AAR | To mutagenize residues 265 and 266 of MinD from KR to AA | 5' TTA TCC TCC CAA CGC CGC GAG GAA GCC TTT CTT CTC3' |
| MinDKR265EEF | To mutagenize residues 265 and 266 of MinD from KR to EE | 5' GAG AAG AAA GGC TTC CTC GAA GAA CGC TTG TTC GGA GGATAA 3' |
| MinDKR265EER | To mutagenize residues 265 and 266 of MinD from KR to EE | 5' TTA TCC TCC GAA CAA GCG TTC TTC GAG GAA GCC TTTCTT CTC 3' |
| MinDΔ8F | To introduce a stop codon 15 residues from the end of *minD* | 5' GAG AAG AAA GGC TAA CTC AAA CGC 3' |
| MinDΔ8R | To introduce a stop codon 15 residues from the end of *minD* | 5' CGC AAA CTC TTA GCC TTT CTT CTC 3' |
| MinDD154AF | To mutagenize residue 154 of MinD from D to A | 5' CGATGGATTTTGAATTTAGCGTTTGTGACTCCCCGG 3' |
| MinDD154AR | To mutagenize residue 154 of MinD from D to A | 5' CCGGGGAGTCACAAACGCTAAATTCAAAATCCATCG 3' |
| MinCR133AF | To mutagenize residue 133 of MinC from R to A | 5' GCTTTTTCCTTTCCTTCGCTCAGGATAGGCAGCCCC 3' |
| MinCR133AR | To mutagenize residue 133 of MinC from R to A | 5' GGGGCTGCCTATCCTGAGCGAAGGAAAGGAAAAAGC 3' |

### Table S3. Copolymer fractions by MinD and MinC variants

| Reaction Conditions for MinD wild type or variant | MinD in copolymer fraction (%) | MinD(D154A) in copolymer fraction (%) |
|---|---|---|
| - ATP | 22.6 ± 5.5 | 15.9 ± 1.8 |
| + ATP | 21.0 ± 2.7 | 25.3 ± 1.9 |
| MinC - ATP | 28.2 ± 7.4 | 24.2 ± 6.5 |
| MinC + ATP | 52.1 ± 7.3 | 21.8 ± 4.0 |
| MinC(R133A) - ATP | 17.7 ± 3.7 | n.t.[1] |
| MinC(R133A) + ATP | 13.8 ± 12.9 | n.t.[1] |

[1] n.t. indicates not tested.

Fig. S1

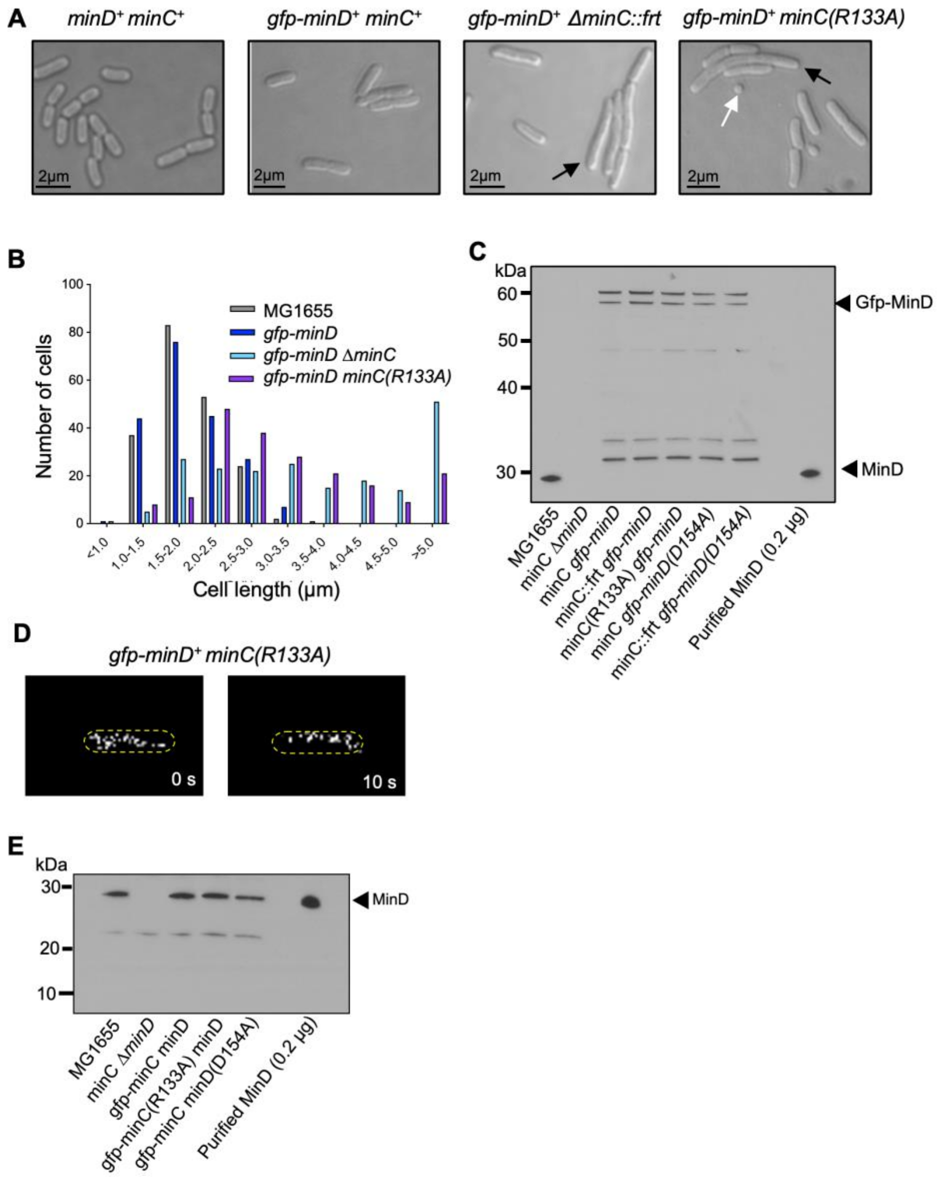


**Fig. S1. MinC(R133A) is defective for interaction with MinD.**
(A) DIC microscopy images of log phase MG1655, MG1655 *gfp-minD* (CF005), MG1655 *gfp-minD ΔminC::frt* (CF315), and MG1655 *gfp-minD minC(R133A)* (CF095) were obtained. Black arrows denote filamentous cells and white arrows denote minicells. Size bar is equal to 2 µm. (B) Cell length distribution of strains shown in (A) indicates that MinC(R133A) (CF095) and Δ*minC* (CF315) have larger distributions of cell length compared to WT cells. (C) Total protein of cell extracts containing MinC-MinD interface mutants, as indicated, in *minD* deletion and *gfp-minD*+ strains were probed with anti-MinD antisera. Western blot images are representative of at least three replicates. (D) Time lapse confocal fluorescence microscopy images of polar Gfp-MinD in a strain expressing Gfp-MinD and MinC(R133A) was captured. (E) Total protein of cell extracts containing MinC-MinD interface mutants, as indicated , in *minD* deletion and *gfp-minD*+ strains were probed with anti-MinD antisera. Western blot images are representative of at least three replicates.

Fig. S2

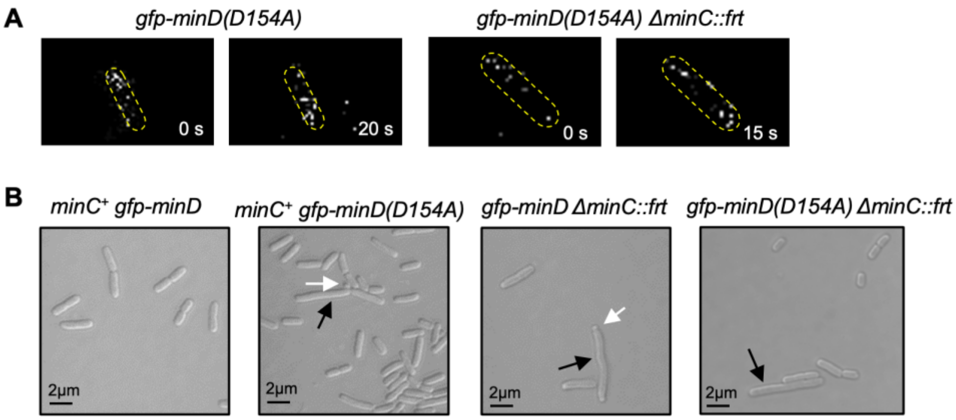


**Fig. S2. MinD(D154A) displays altered morphology and oscillates *in vivo*.**
(A) Time lapse confocal fluorescence images of Gfp-MinD at the poles during one period in strains harboring Gfp-MinD(D154A) with (CF089) and without MinC (CF313). (B) To compare morphology of the MinD(D154A) variants, we visualized log phase cells by DIC microscopy, including MG1655 *gfp-minD* (CF005), MG1655 *gfp-minD(D154A)* (CF089), MG1655 *gfp-minD ΔminC::frt* (CF315), and MG1655 *gfp-minD(D154A) ΔminC::frt* (CF313). Black arrows denote filamentous cells, white arrows denote minicells. Scale bar is 2 μm.

Fig. S3

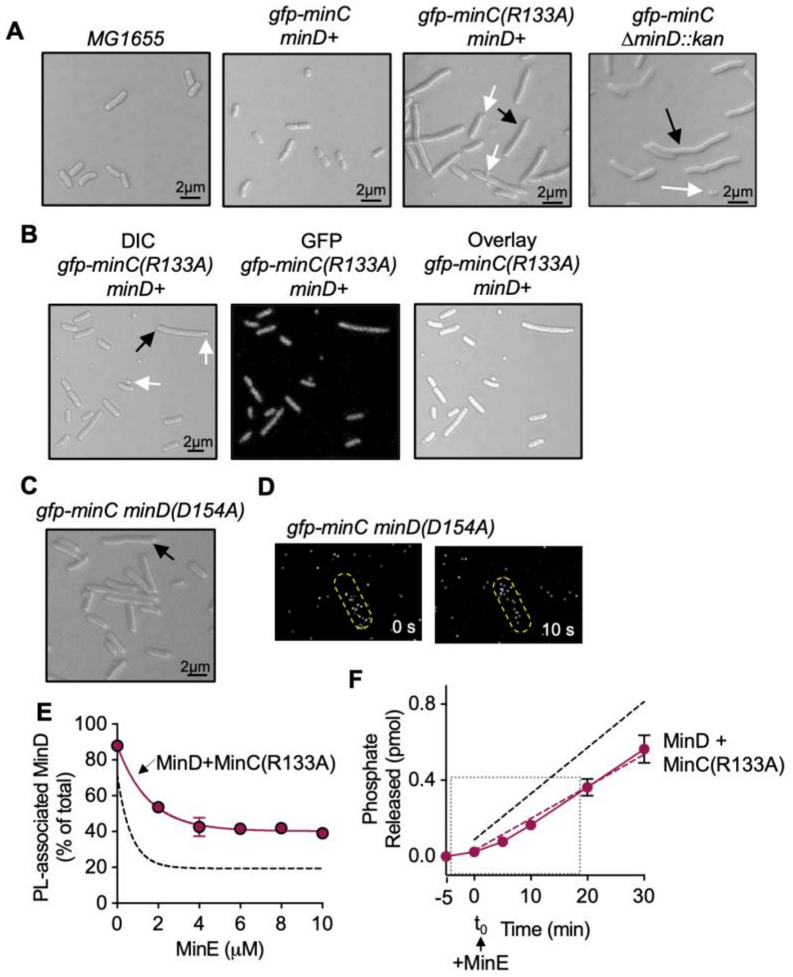


**Fig. S3. MinD-MinC interaction site variants confer altered cell morphology and oscillation.**
(A) Morphologies of log phase cells containing *gfp-minC(R133A)* or deleted for *minC* were compared by DIC microscopy of MG1655,, MG1655 *gfp-minC* (CL0248), MG1655 *gfp-minC(R133A*) (CF266), and MG1655 *gfp-minC ΔminD::kan* (CF0481). Black arrows denote filamentous cells while white arrows denote minicells. Scale bar is 2 µm. (B) To visualize diffuse fluorescence, an overlay of fluorescence and DIC microscopy of MG1655 *gfp-minC(R133A)* (CF266) is shown. Black arrows denote filamentous cells while white arrows denote minicells. Scale bar is 2 µm. (C) Log phase MG1655 *gfp-minC minD(D154A)* cells containing *gfp-minC* and *minD(154A)* were imaged by DIC microscopy (CF086). Black arrows denote filamentous cells while white arrows denote minicells. Scale bar is 2 µm. (D) Time lapse fluorescence images of Gfp-MinC localization in a strain containing MinD(D154A) are shown (CF086). (E) To evaluate MinE-stimulated release, we plotted the percent of MinD that remained associated with SUVs after incubating with an increasing concentration of MinE 0 µM to 10 µM). Dashed line shows the release curve generated by incubating MinD with MinC and increasing MinE, as shown in Figure 4D for comparison. (F) Staged ATP hydrolysis assays were performed where MinD (8 µM) and MinC(R133A) (4 µM) are preincubated with ATP (4mM), and phospholipids (250 µg ml$^{-1}$) for 5 minutes at 30 °C, when MinE (16 µM) is then added and total phosphate released is monitored for 30 minutes. Dashed line shows linear ATP hydrolysis detected by incubating MinD with MinC and then adding MinE, as shown in Figure 4E for comparison. Data shown in panels E and F are representative of at least three replicates. Error is shown as standard error.

Fig. S4

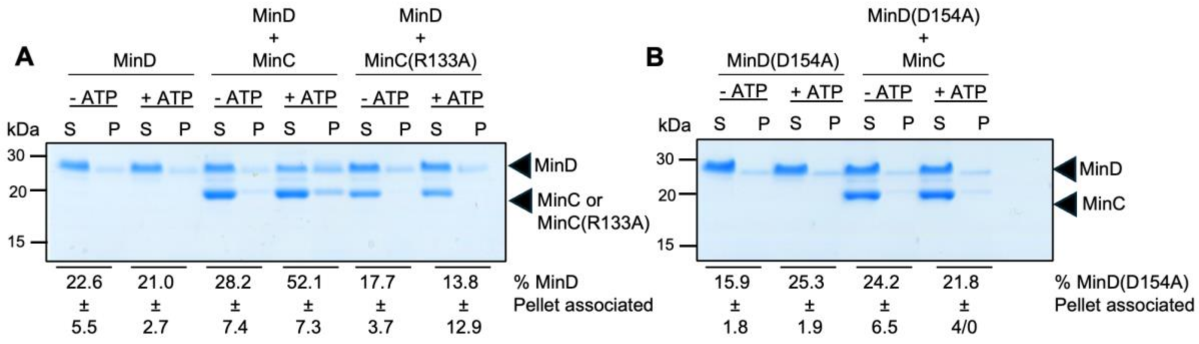


**Fig. S4. MinC-MinD binding site variants are defective for copolymer formation in vitro.** (A) An ultracentrifugation assay of MinD (8 μM) with and without ATP (4 mM), with and without MinC or MinC(R133A) (4 μM) was performed. Copolymer formation was evaluated by centrifugation and SDS-PAGE of soluble (S) and protein pellet (P) fractions, and fractions were quantified by Coomassie staining and densitometry. (B) An ultracentrifugation assay of MinD(D154A) (8 μM) with and without ATP (4 mM), with and without MinC (4 μM) was performed as in (A). In panels A-B, gel images are representative of at least three replicates. Error is shown as standard error.

Fig. S5

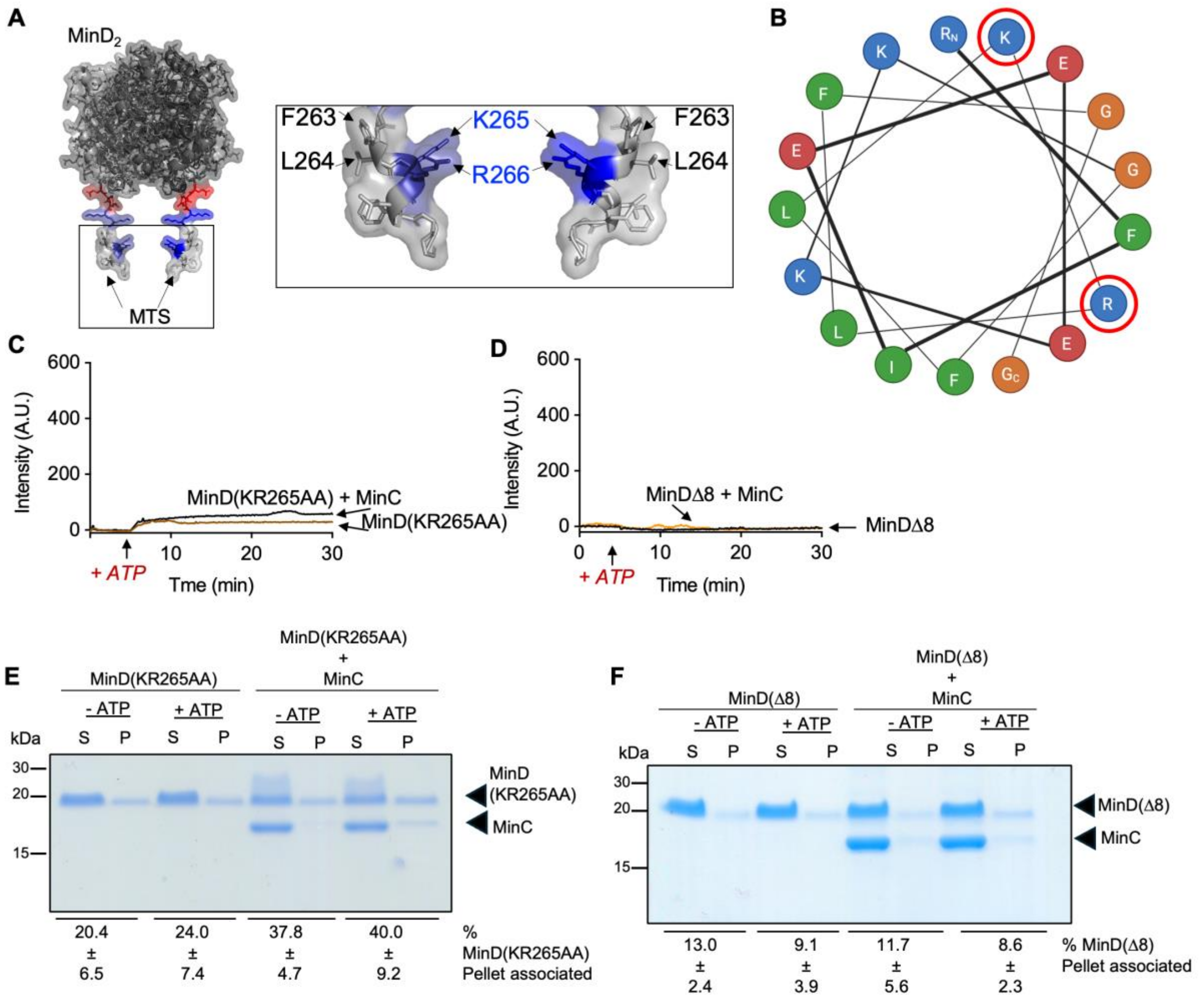


**Fig. S5. MinD(KR265AA) and MinD(Δ8) are defective for formation of copolymers in vitro.** (A) The crystal structure of the MinD dimer (PDB 3Q9L) was modeled with highlighted MTS residues(18). Red residues denote the beginning of the MTS, while blue indicate the residues K265 and R266. (B) A representation of the amphipathic helix wheel of wild-type MinD, including the last 17 residues of MinD is shown. Residues K265 and R266 are circled in red. (C) Light scattering (90°) of MinD(KR265AA) (8 μM), alone and with MinC (4 μM), with ATP (8 mM) added at 5 minutes. (D) Light scattering of MinDΔ8 (8 μM) as in (C). (E) Ultracentrifugation assay of MinD(KR265AA) (8 μM) with and without ATP (8 mM), with and without MinC (4 μM). Copolymer formation was evaluated by centrifugation and SDS-PAGE of soluble (S) and protein pellet (P) fractions, and fractions were quantified by Coomassie staining and densitometry. (F) Ultracentrifugation assay of MinD(KR265AA) (8 μM) as in (E). Data shown in C-F is representative of at least three replicates.

Fig. S6

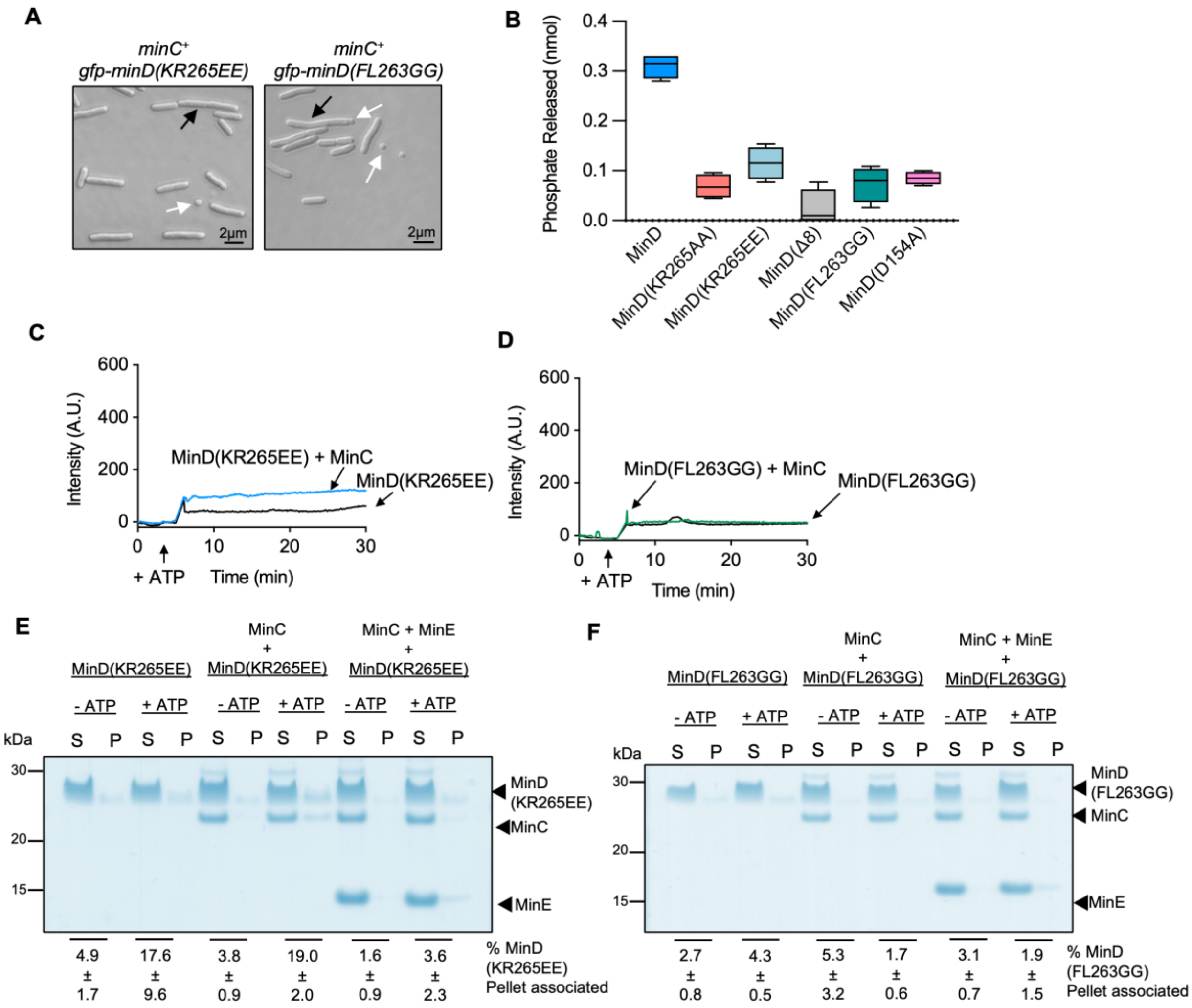


**Fig. S6. MinD(KR265EE) and MinD(FL263GG) are defective for function**
(A) Log phase MG1655 *gfp-minD(KR265EE)* (CF255) and MG1655 *gfp-minD(FL263GG)* (CF150) cells were visualized by DIC microscopy. Black arrows denote filamentous cells while white arrows denote minicells. Scale bar is 2 µm. (B) ATP hydrolysis of MinD MTS variants was measured endpoint as indicated in reaction containing SUVs and MinE. (C) Light scattering of MinD(KR265EE) (8 µM) alone or with MinC (4 µM), with ATP (8 mM) added at 5 minutes. (D) Light scattering of MinD(FL263GG) (8 µM) as in (C). (E) Phospholipid binding assay of MinD(KR265EE) with and without ATP and MinC as indicated. Release by MinE was evaluated by centrifugation and SDS-PAGE of soluble (S) and phospholipid-associated (P) fractions. Gel is a representative image of at least three replicates. (F) Phospholipid binding assay of MinD(FL263GG) with and without ATP, with and without MinC. Release was evaluated by centrifugation and SDS-PAGE of soluble (S) and phospholipid-associated (P) fractions and fractions were quantified by Coomassie staining and densitometry. Data shown is representative of at least three replicates.

Fig. S7

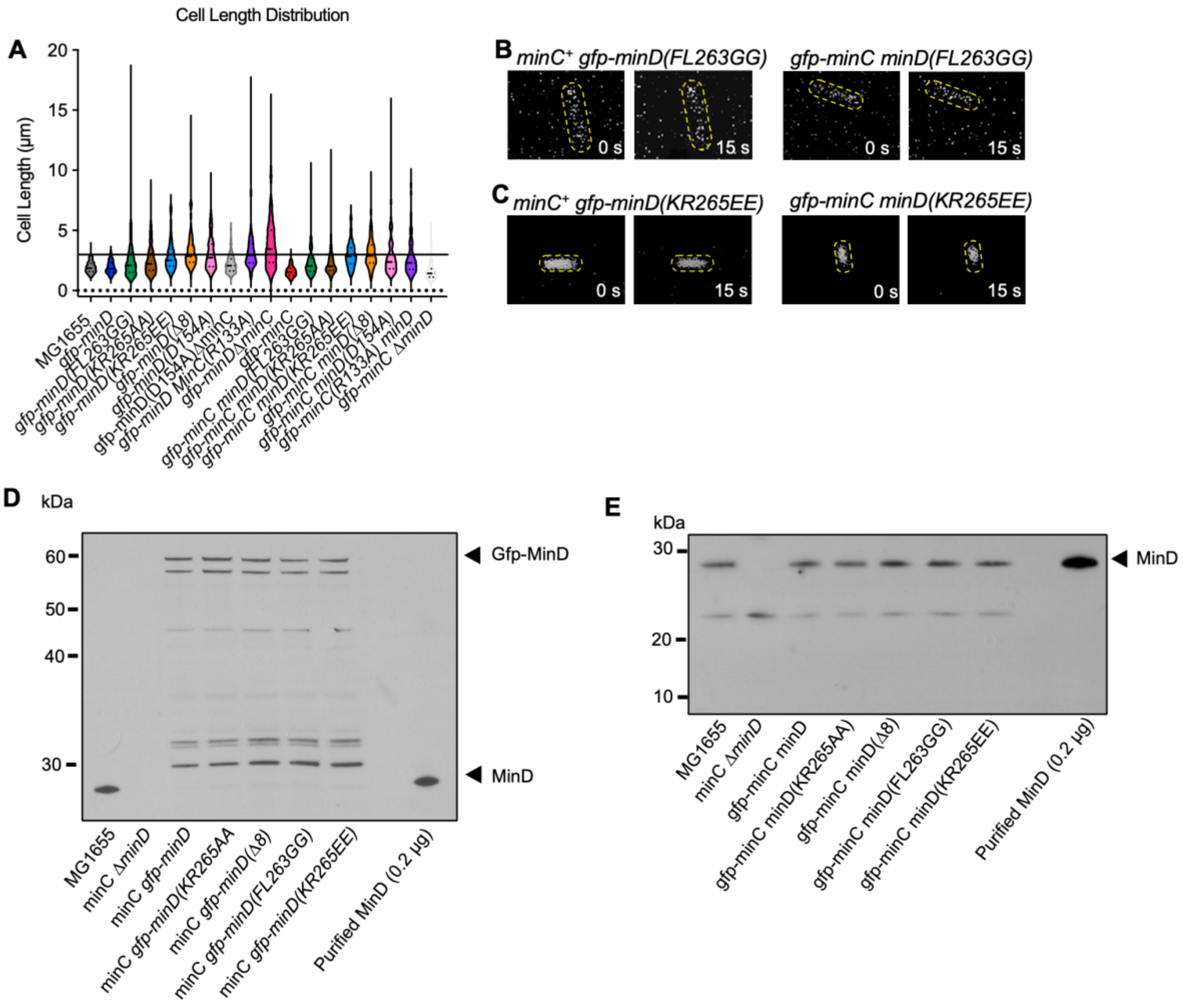


**Fig. S7. Mutations in the membrane targeting sequence (MTS) of MinD impair oscillation in vivo.**

(A) A cell length distribution plot of strains containing *minD(FL263GG)* (CF150, CF254), *minD(KR265AA)* (CF097, CF071), *minD(KR265EE)* (CF255, CL0482), and *minD(Δ8)* (CF256, CF070) in both *gfp-minD* and *gfp-minC* backgrounds is shown for comparison to a WT strain (MG1655). At least 200 cells were measured for every strain. (B) Strains expressing Gfp-MinD(FL263GG) (CF150) or MinD(KR265EE) (CF255) did not exhibit oscillation by confocal fluorescence microscopy; diffuse cytoplasmic fluorescence was observed. (C) Gfp-MinC did not oscillate in strains expressing MinD(FL263GG), and MinD(KR265EE) and diffuse cytoplasmic fluorescence was observed. (D) Total protein of cell extracts containing MinD MTS variants in the *gfp-minC* background was probed with anti-MinD sera for comparison with wild type MinD expression. Purified MinD (0.2 µg) was added as a loading control. (E) Total protein of cell extracts containing MinD MTS variants in the *gfp-minC* background was probed with anti-MinD sera. Purified MinD (0.2 µg) was added as a loading control. Western blot images in (D) and (E) are representative of at least three replicates.
